# T:B cell cooperation in ectopic lymphoid follicles propagates CNS autoimmunity

**DOI:** 10.1101/2023.06.01.543004

**Authors:** Anna Kolz, Isabel Julia Bauer, Clara de la Rosa, Vladyslav Kavaka, Anna Sophie Thomann, Rosa Schmitz, Martin Kerschensteiner, Eduardo Beltran, Naoto Kawakami, Anneli Peters

## Abstract

Meningeal ectopic lymphoid follicle-like structures (eLFs) have been described in multiple sclerosis (MS) and its animal model experimental autoimmune encephalomyelitis (EAE), but their role in CNS autoimmunity is unclear. To analyze the cellular phenotypes and interactions within these structures, we employed a Th17 adoptive transfer EAE model featuring formation of large, numerous eLFs. Single-cell transcriptomic analysis revealed that clusters of activated B cells and B1/Marginal Zone-like B cells are overrepresented in the CNS and identified B cells poised for undergoing antigen-driven germinal center (GC) reactions and clonal expansion in the CNS. Furthermore, CNS B cells showed enhanced capacity for antigen presentation and immunological synapse formation compared to peripheral B cells. To directly visualize Th17:B cell cooperation in eLFs, we labeled Th17 cells with a ratiometric calcium sensor, and tracked their interactions with tdTomato-labeled B cells in real-time. Thereby, we demonstrated for the first time that T and B cells form long-lasting antigen-specific contacts in meningeal eLFs that result in reactivation of autoreactive T cells. Consistent with these findings, autoreactive T cells depended on CNS B cells to maintain a pro-inflammatory cytokine profile in the CNS. Collectively, our study reveals that extensive T:B cell cooperation occurs in meningeal eLFs in our model promoting differentiation and clonal expansion of B cells, as well as reactivation of CNS T cells and thereby supporting smoldering inflammatory processes within the CNS compartment. Our results provide valuable insights into the function of eLFs and may provide a direction for future research in MS.

## Introduction

In multiple sclerosis (MS) autoreactive T cells recognizing central nervous system (CNS)-specific antigens are thought to be important drivers of disease. To our current knowledge, they are primed in the periphery before they invade the CNS, where they presumably become reactivated by contact with the autoantigen, and subsequently orchestrate the inflammatory cascade. Both Th1 and Th17 cells contribute to pathogenesis in MS and its animal model experimental autoimmune encephalomyelitis (EAE), and during the past two decades, compelling evidence has also suggested a pivotal role of B cells (Comi et al., 2021). However, it is not clear how T cells and B cells cooperate to induce and propagate disease. In MS, ectopic lymphoid follicle-like structures (eLFs) were found in the leptomeninges of patients with progressive disease course (Magliozzi et al., 2007; Serafini et al., 2004), but also in earlier stages of relapsing-remitting MS (Lucchinetti et al., 2011). Initially, they were thought to be restricted to brain meninges, where they are localized mainly in sulci, but a recent study showed that they can also form in spinal cord meninges (Reali et al., 2020). These aggregates show varying degrees of maturation, ranging from loose clusters of cells to highly organized follicle structures, which contain plasma cells, proliferating B cells, T cells and follicular dendritic cells (FDCs), and therefore are suggestive of germinal center formation (Mitsdoerffer and Peters, 2016). Clinical data show a correlation between the presence of eLFs and a more severe disease course – younger age at disease onset, irreversible disability, and death, as well as a more pronounced cortical pathology in the tissue underneath eLFs (Howell et al., 2011; Magliozzi *et al*., 2007; Magliozzi et al., 2010; Serafini et al., 2007; Serafini *et al*., 2004). Therefore, these follicles have been postulated to maintain or exacerbate chronic inflammation directly in the CNS. Pathogenic functions of eLFs could include both initial priming and maturation of autoreactive B and T cell clones, as well as reactivation and expansion of pre-existing autoreactive T and B cells recruited from the periphery. In addition, eLFs may be a source of pathogenic soluble factors such as auto-antibodies and pro-inflammatory cytokines that affect the underlying regions. However, eLFs might also harbor regulatory processes aimed at dampening excessive inflammation and it is possible that both eLFs with disease promoting and eLFs with regulatory function exist in MS either in different subsets of patients or in different phases of the disease. Since data from human eLFs is based on and largely restricted to histological analysis of post-mortem CNS material, our knowledge of cellular processes within eLFs remains limited.

To study eLFs in more detail, here we employed a Th17 adoptive transfer EAE model, which features formation of eLFs within the meninges (Jager et al., 2009; Peters et al., 2011). While meningeal eLFs were also identified in other EAE models differing regarding genetic background, mode of EAE induction and auto-antigen (Dang et al., 2015; Kuerten et al., 2012; Magliozzi et al., 2004; Pikor et al., 2015), we chose the Th17 adoptive transfer EAE model for our study as it exhibits robust formation of the largest and most numerous eLFs. Due to the relatively short and severe disease course, the Th17 EAE model is not suitable to study the clinical impact of eLFs, however, it can provide valuable insights into the early stages of eLF formation. Specifically, the large number and size of the eLFs and the substantial recruitment of B cells facilitate in depth phenotyping via single-cell RNA sequencing and even enable us to study cellular interactions via live imaging.

The aim of this study was to investigate the unique relationship of Th17 cells and B cells in EAE development, and, in particular, to assess the nature of their interaction in eLFs. Combining histological and flow cytometric analyses with single-cell transcriptomics and intravital imaging, we show that meningeal eLFs are sites of extensive T:B cell cooperation resulting in both development of germinal center B cells and reactivation of autoreactive T cells and can thereby perpetuate autoimmune reactions in the CNS compartment.

## Results

### Th17 cells induce formation of ectopic lymphoid follicles in CNS meninges

To be able to study cellular processes in eLFs, we first established adoptive transfer EAE according to Jäger et al. (Jager *et al*., 2009), a model in which disease is induced by transfer of *in vitro* differentiated pure MOG-specific Th1 or Th17 cells into immunocompetent recipient mice (Figure 1A, S1A). As previously shown, Th1 and Th17 cells induced EAE independently of each other with comparable clinical disease courses and similar severity (Figure S1B). In the inflamed CNS, transferred Th1 and Th17 cells maintained stable expression of their signature cytokine IFNγ and IL-17A, respectively, and Th17 cells can additionally adopt IFNγ production (Figure S1C, D). To assess B cell recruitment to the CNS, we compared B cell frequencies among CNS-infiltrating cells between Th1 and Th17 recipient mice at late peak of disease. We detected a significant increase in both frequency and absolute numbers of B cells in the CNS of Th17 recipients compared to Th1 recipients (Figure 1B-D). We next performed a detailed immunohistochemical analysis of CNS lesions to investigate the localization, structural organization, and composition of cellular infiltrates in Th17-EAE mice. Following identification of regions with lymphocytic infiltrates by Giemsa staining (Figure S2A), neighboring sections were stained for T cells, B cells and laminin, which visualizes the basal lamina of blood vessels and the pia mater. In line with previous observations in Th17 recipient mice (Peters *et al*., 2011; Pikor *et al*., 2015), we found numerous eLFs within the meninges of the spinal cord. eLFs consisted of accumulated T and B cells and showed varying degrees of organization and size: while some contained loose aggregates of a few cells, others were composed of large amounts of T and B cells and extended over wide distances of up to several millimeters (Figure 1E, S2B). Based on their unique morphology and expression of CD11b, we also identified macrophages in close vicinity, but also within the follicle structures (Figure S2D). Large eLFs were also found in many different areas of the brain, including for example the subarachnoid space at the base of the brain (Figure S2C) and the cortex (Figure 1F). Similarly, we did not observe a preferential location for eLFs within spinal meninges but rather a widespread distribution along the cord (Figure 1G). eLFs appeared to originate from the meninges and invade the underlying parenchyma, and often exhibited separate T and B cell zones in our model, thereby mimicking the structural organization of SLOs. Notably, all eLFs were characterized by prominent B cell clustering, with B cells being almost exclusively confined to these meningeal aggregates. In contrast, T cells were more widely distributed and detected both inside eLFs in large numbers, as well as in the underlying subpial parenchymal tissues (Figure 1E, F, S2B, C). Similar to eLFs in human CNS tissue (Magliozzi *et al*., 2007), staining for Ki67 revealed eLFs in our model to be sites of T and B cell proliferation (Figure 1H). To evaluate the degree of eLF maturation, we assessed expression of GC markers on B and T cells. In many eLFs, only few scattered B cells expressed the GC markers PNA or AID (Figure 1I, J), however, some eLFs exhibited clusters of AID^+^ B cells (Figure 1K), which have also been detected in human eLFs (Serafini *et al*., 2007). In addition, PNA^+^ B cells (Figure S3A), isotype-switched IgM^-^ IgD^-^ B cells (Figure S3B), and a small population of plasma cells (Figure S3C) were detected among CNS-infiltrating cells via flow cytometry. Furthermore, a fraction of CNS-infiltrating T cells expressed the GC and activation marker GL7 with transferred 2D2 Th17 cells showing higher frequency of GL7^+^ cells than endogenous T cells from the recipient (Figure S3D).

**Figure 1.**
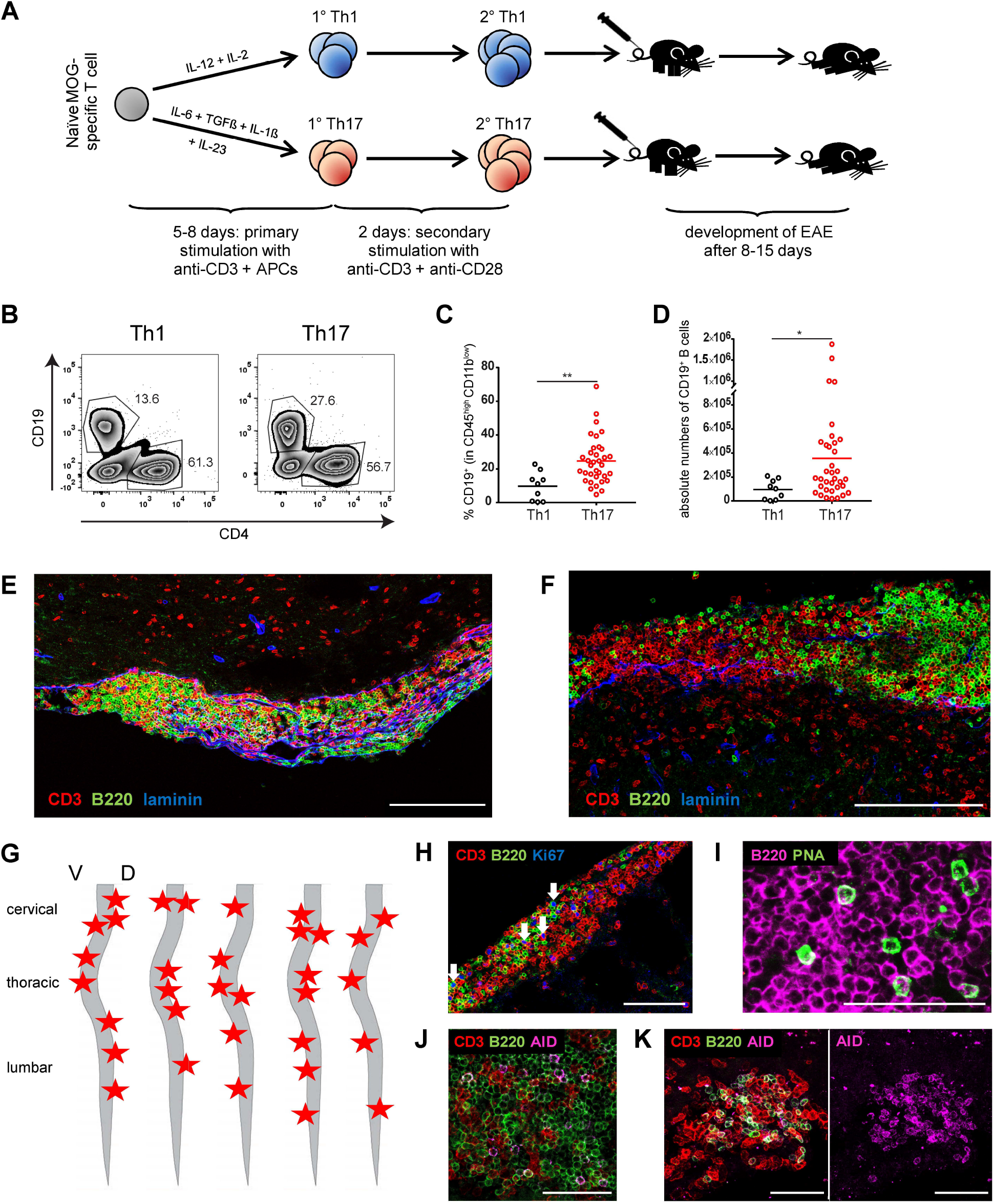
Th17 cells induce formation of large ectopic lymphoid follicle-like structures expressing germinal center markers in association with the meninges of spinal cord and brain. (**A**) Experimental outline. Naïve MOG-specific T cells isolated from 2D2 mice are differentiated with polarizing cytokines into Th1 or Th17 cells in the presence of anti-CD3 and APCs. After 5-8 days, the cells are restimulated with anti-CD3 and anti-CD28 for 2 days before they are transferred i.v. or i.p. into recipient animals. Clinical disease develops after 8-15 days post transfer. (**B-D**) At the peak of disease, cells were isolated from the CNS of Th1 and Th17-EAE mice and analyzed for the presence of CD19^+^ B cells. (**B**) Representative flow cytometry plots (pre-gated on CD45^high^ CD11b^low^ cells) and quantification of the (**C**) frequency of CD19^+^ B cells in CD45^high^ CD11b^low^ cells and (**D**) absolute numbers of infiltrating CD19^+^ B cells in the CNS. Graphs show cumulative data from three (Th1-EAE) or eight (Th17-EAE) independent experiments. Mann-Whitney U test. *p < 0.05; **p < 0.01. Horizontal lines indicate means. Dots represent individual mice. (**E, F**) At the peak of disease, (**E**) longitudinal spinal cord and (**F**) coronal brain cryosections of Th17-EAE mice were prepared and stained for T cells (CD3, red), B cells (B220, green) and laminin (blue). (**E**) Zoom of red highlighted area in Figure S2B. (**F**) Cortex. Scale bars: 200 µm. (**G**) Scheme shows the spatial distribution of the largest ectopic lymphoid aggregates (red stars) in the spinal cord of the five mice analyzed. *V, ventral; D, dorsal.* (**H-K**) CNS cryosections of Th17-EAE mice at the peak of disease were stained for (**H**) T cells (CD3, red), B cells (B220, green) and the proliferation marker Ki67 (blue) (white arrows indicate Ki67^+^ T and B cells); (**I**) B cells (B220, magenta) and the germinal center marker peanut agglutinin (PNA, green); (**J**, **K**) T cells (CD3, red), B cells (B220, green) and the enzyme activation-induced cytidine deaminase (AID, magenta). For better visualization, the AID staining is shown separately in K. Scale bars: 50 µm.

Taken together, we confirmed that the Th17 adoptive transfer EAE model prominently features formation of eLFs reminiscent of eLFs in MS patients in both spinal cord and brain meninges. Considering the high frequency of recruited B cells, the high abundance and large size of eLFs, as well as our initial evidence that at least some of the eLFs are mature enough to harbor GC B and T cells, our model is well suited to study the cellular processes in CNS eLFs in more detail.

### CNS-infiltrating B cells are highly activated and poised for undergoing germinal center reactions

To obtain a more refined understanding of the properties and effector functions of B cells in meningeal eLFs, we characterized the transcriptome of CNS-infiltrating B cells from Th17 recipients by single-cell RNA sequencing, and compared it to the transcriptome of peripheral B cells from spleen and cervical lymph nodes (cLN) of the same animal. Unbiased clustering of the single-cell transcriptomes revealed ten individual B cell clusters varying substantially in size and containing B cells of all three compartments. Interestingly, CNS B cells showed a distinct distribution of B cells across the different clusters, indicating them to be phenotypically different from peripheral B cells (Figure 2A, Figure S4A+B). Among the identified clusters, cluster 6-9 were highly enriched for CNS B cells, most pronounced for cluster 6, in which 93.5 % of cells belonged to the CNS fraction (Figure 2B). Gene set enrichment analysis against the GO biological processes database revealed that cluster 6 B cells are metabolically highly active and have a high energy demand as they showed upregulation of pathways associated with respiration, ATP biosynthesis, glycolysis, fatty acid metabolism and oxidative phosphorylation (Figure S4C, S5A). In line with this, they showed highly increased expression of many ribosomal proteins (Figure S4A and Table S1), which have been shown to be upregulated in early GC B cells in response to mTOR signaling in order to prepare for the increased translational demand of an imminent GC reaction (Ersching et al., 2017). Notably, both mTOR and Myc signaling, which precede entry into a GC reaction, were high in cluster 9, 6 and 7 B cells, and always higher in CNS B cells compared to peripheral B cells within each cluster (Fig. S4C). Pathway analysis showed that cluster 7 B cells are highly activated / differentiated and transcriptionally very active (Fig. S5B). Thus, *Nr4a1*, *Egr1, Egr2* and *Egr3*, which are promptly induced upon B cell activation through the BCR, are among the top 10 up-regulated genes in cluster 7 (Fig. S4A and Table S1). Cluster 9 cells showed upregulation of pathways associated with cell cycle, mitosis, DNA replication, and proliferation and accordingly *Mki67* was among the top 10 upregulated genes in this cluster (Fig. S5C, Fig. S4A and Table S1). Importantly, the CNS-enriched clusters 6 and 7 showed highly significantly increased expression of genes associated with a light zone (LZ) germinal center B cell signature (Victora et al., 2010), compared to other CNS B cells as well as compared to peripheral B cells within the same cluster (Figure 2C). Thus, for example the LZ GC markers *Cd83* and *Fas*, as well as the cyclin *Ccnd2* and the transcription factor *Bhlhe40*, which have been associated with LZ GCs (Rauschmeier et al., 2022; Victora *et al*., 2010), were among the top 50 upregulated genes in both cluster 6 and 7 (Table S1). These data indicate that the phenotype of B cells especially in cluster 6 and 7, which are highly overrepresented in the CNS, is compatible with LZ GC B cells, suggesting that a considerable fraction (about 19.5 %) of CNS B cells is poised to undergo GC reactions. In contrast, only cluster 9 B cells showed highly elevated expression of genes associated with dark zone (DZ) germinal center B cells compared to other CNS B cells (Victora *et al*., 2010) (Figure 2D), including for example *Stmn1, Cenpa,* and the cyclin *Ccnb2*. This is in line with the increased proliferative capacity of these cells indicated by the pathway analysis (Fig. S5C) and the high level of oxidative phosphorylation (Fig. S4C), suggesting that with cluster 9 only a very small fraction (1.4 %) of B cells in eLFs has progressed to DZ GC B cells to begin clonal expansion.

**Figure 2.**
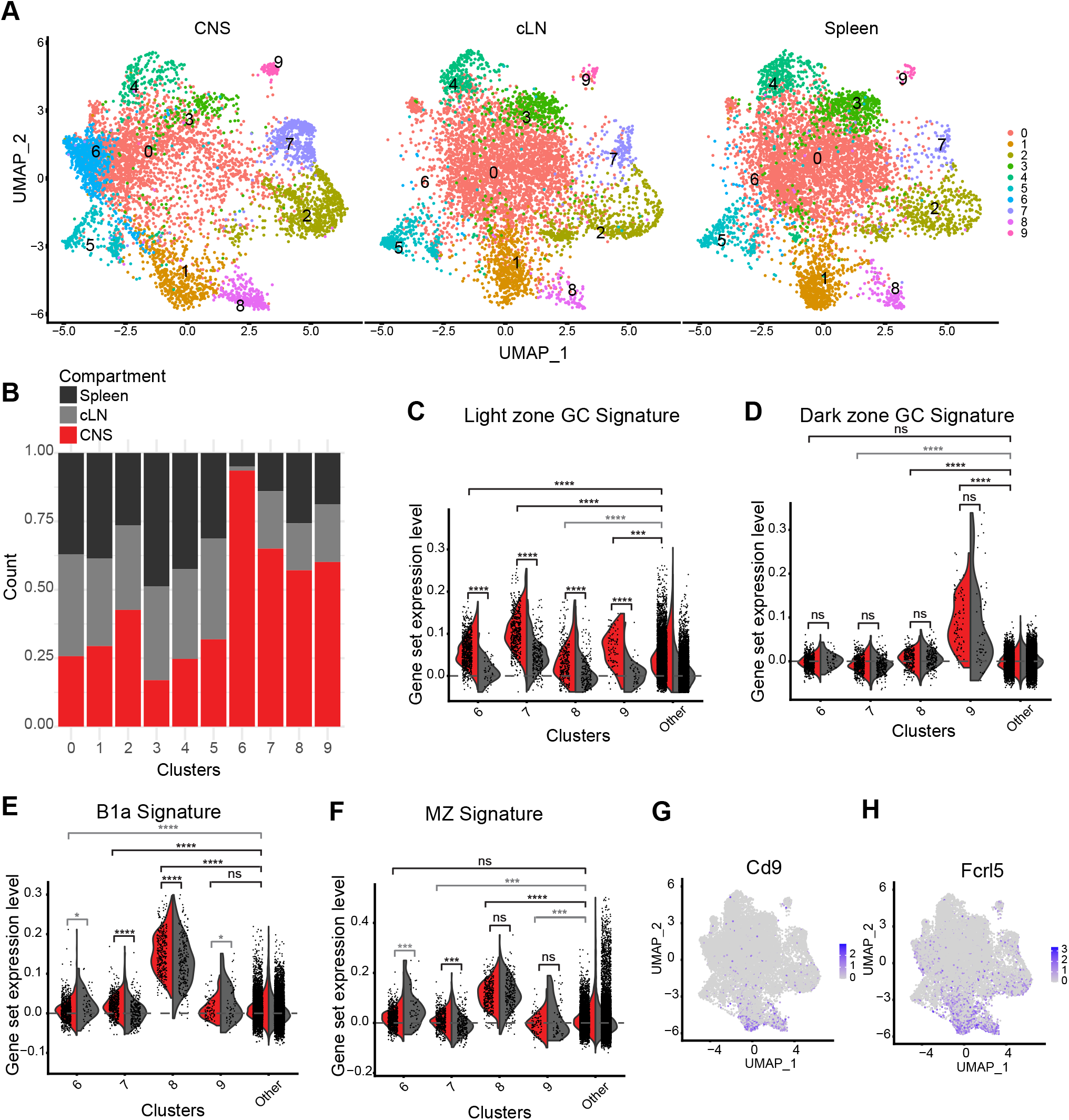
CNS-infiltrating B cells are phenotypically different from peripheral B cells. At the peak of disease, B cells were sorted from indicated organs of Th17 recipients (n = 3 mice) based on their expression of CD19, and their gene expression was characterized by single-cell RNA sequencing. (**A**) Single cell transcriptomes are depicted in UMAP plots representing ten color-coded cell clusters, shown separately for CNS, cervical lymph node (cLN) and spleen. (**B**) For each cluster, the fraction of cells belonging to CNS, cLN or spleen was calculated. (**C, D**) Violin plots showing expression levels of gene sets associated with (**C**) light zone and (**D**) dark zone germinal center (GC) B cells. (**E, F**) Violin plots showing expression levels of gene sets associated with (**E**) B1a B cells and (**F**) MZ B cells. (**G**, **H**) Feature plots depicting distribution of (**G**) Cd9 or (**H**) Fcrl5-expressing B cells among B cell clusters. Two-sided *t*-test with FDR adjusted p-values. *p < 0.05; **p < 0.01; ***p < 0.001, ****p < 0.0001. Significantly downregulated gene sets are marked in grey.

### B1a / MZ-like B cells infiltrate the CNS

In addition to the clusters described above, also cluster 8 was highly enriched for CNS B cells. Further examination of the upregulated genes in this cluster, such as *Mzb1*, *Fcrl5* and *Cd9* suggested similarities to marginal zone (MZ) and B1 cells (Fig. S4A, Table S1). This observation was validated by comparing the cluster 8 transcriptome with publicly available gene signatures of B1a cells from spleen and peritoneum and splenic MZ B cells confirming that cluster 8 represents a population of B1a / MZ-like B cells in the CNS (Figure 2E, F). The only other cluster that showed highly elevated expression of B1a and MZ markers including *Cd9* and *Fcrl5* was cluster 1, which consists mainly of peripheral B cells and contains the MZ B cells in the spleen (Fig. 2G, H, Fig. S4A, Table S1). As regular MZ B cells are often identified via expression of CD1d and CD21, we analyzed CNS-infiltrating cells for the presence of IgM^high^ IgD^low^ CD21^high^ CD1d^+^ B cells via flow cytometry, and detected a small and relatively dim population (Figure S5D, E). Additionally, we confirmed presence of scattered CD1d^+^ B cells in eLFs in the CNS via confocal microscopy (Figure S5F). Considering that CD1d is not necessarily expressed on B1a cells, this gating and staining strategy may not have included all the cells in cluster 8.

### CNS-infiltrating B cells undergo clonal expansion

Analysis of the BCR repertoire via comparison of the sequence of Ig variable regions revealed the existence of expanded B cell clones, and interestingly, the clones with the highest expansion factors predominantly belonged to cluster 8, especially in the CNS (Figure 3A, S5G, H). Importantly, 55.5 % of expanded cells represented B cells from the CNS, while 28.8 % and 15.5 % were derived from spleen and cLN, respectively (Figure 3B). This preferential localization of expanded clones may indicate that infiltrating B cells undergo clonal expansion in the inflamed CNS. The clones with the highest expansion factors in the CNS were predominantly localized in cluster 8, but we also found clones with lower expansion factors in the CNS in cluster 9 and 4 (Figure 3C). Since B cells were sorted based on their expression of CD19, plasma cells, which no longer express CD19, were not included in this analysis. Therefore, the expanded CNS B cells could be in the process of differentiating into plasmablasts. Indeed, only cluster 8 and cluster 9 B cells showed significantly increased expression of plasmablast-signature genes compared to other CNS B cells (Figure 3D). Considering that cluster 9 B cells in the CNS displayed features of DZ GC B cells (Fig. 2D), these data suggest that cluster 9 B cells are starting to undergo clonal expansion with still low expansion factors and are also upregulating the plasmablast differentiation program at the beginning of an antigen-driven GC reaction. In contrast, cluster 8 B cells have much higher expansion factors but show lower expression of GC markers, indicating that they may differentiate into plasmablasts independent of classical T cell-driven GC reactions. This is in line with the known properties of regular B1 and MZ B cells to be able to operate more independently of T cell help and give rise to extrafollicular plasmablasts (Cyster and Allen, 2019). A comparison between the transcriptome of non-expanded and expanded cells in cluster 8 revealed that the expanded cells upregulated genes associated with plasma cell differentiation and MZ B cells or B1 B cells, including *Fam46c*, *Emb* and *Gpx4* (Figure S5J). Additionally, cluster 8 B cells showed upregulation of pathways associated with regulation of immune responses (Figure S5I) and expressed for example *Ctla4* and *Apoe* among the top 50 upregulated genes (Fig. S4A, Table S1).

**Figure 3.**
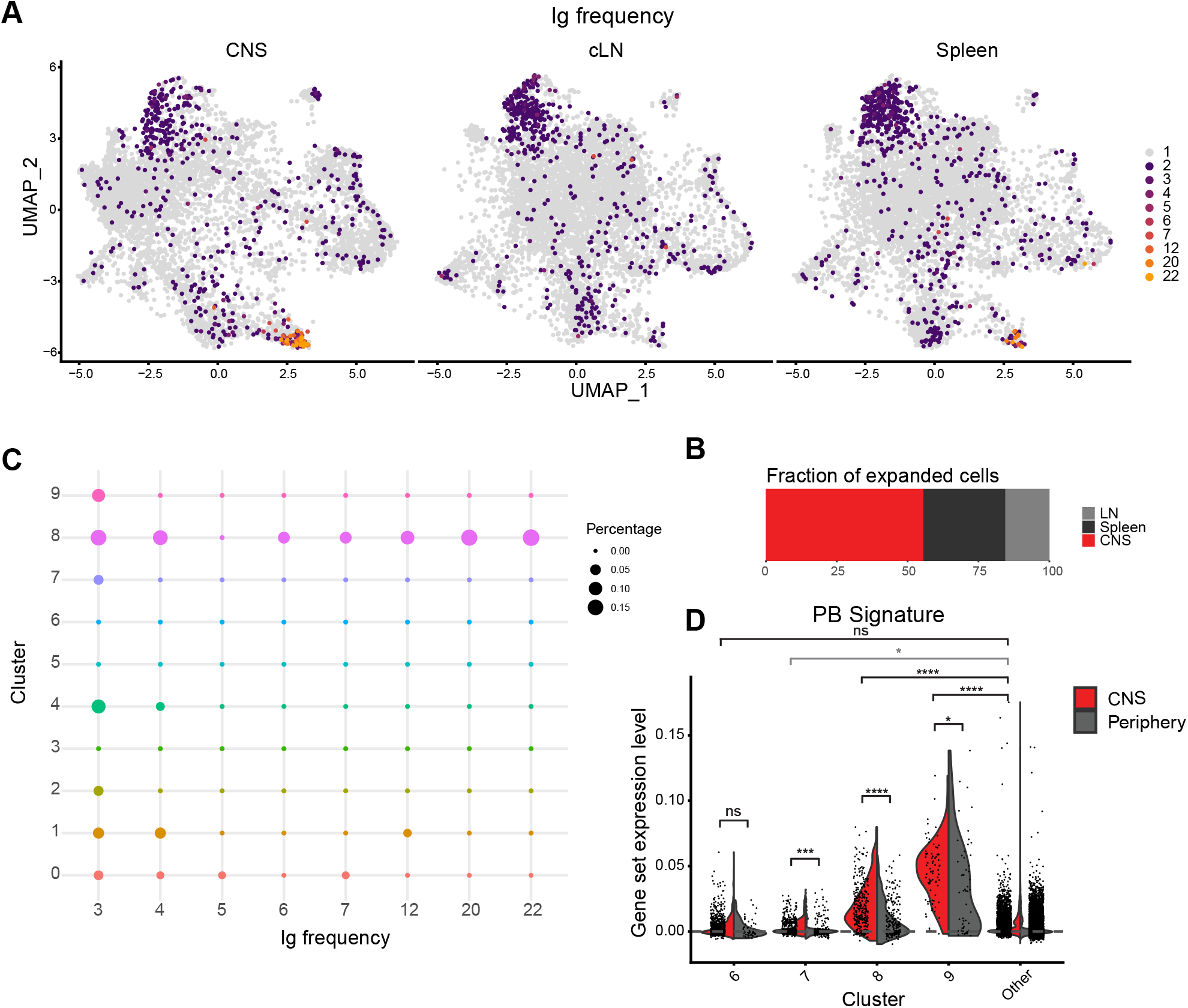
CNS-infiltrating B cells undergo clonal expansion in the CNS. (**A**) At the peak of disease, B cells were sorted from indicated organs of Th17 recipients (n = 3 mice) based on their expression of CD19, and characterized by BCR repertoire analysis. Feature plots depicting distribution of nonexpanded (grey) and expanded (colored) B cells among B cell clusters, plotted separately for CNS, cLN and spleen. The color code indicates the number of cells per clone. (**B**) Fraction of expanded cells belonging to CNS, spleen and cLN. (**C**) Percentage of clones with expansion factors ≥3 for each cluster in the CNS. (**D**) Violin plots showing expression levels of genes associated with plasmablasts (PB). Two-sided *t*-test with FDR adjusted p-values. *p < 0.05; **p < 0.01; ***p < 0.001, ****p < 0.0001.

As we observed that the CNS in Th17 recipient mice supports clonal expansion of B cells and harbors B cells poised to undergo GC reactions, we next investigated whether development and expansion of MOG-specific B cells from the polyclonal repertoire occurs in our model. We first determined the presence of a MOG-specific IgG1 response in the serum of Th17 and Th1 recipients via ELISA. At late peak of clinical disease, many Th17 recipients showed high levels of MOG-specific IgG1 antibodies compared to very low levels before EAE onset. Some Th1 recipients also exhibited elevated antibody titers at the peak of disease compared to very low levels in mice, which were analyzed at an early disease stage before B cells are starting to infiltrate into the CNS. However, on average MOG-specific IgG1 titers in Th17 recipients were about 3-fold higher than in Th1 recipients (Figure S6A). To determine whether CNS eLFs contain MOG-specific B cells, we optimized detection of MOG-specific B cells via IHC by staining CNS cryosections with a fluorescently labeled tetramerized MOG- protein (MOG_tet_), which was validated on LN cryosections from IgH^MOG^ mice carrying MOG-specific B cells (Figure S6B). Staining of CNS cryosections from Th17-EAE mice revealed that most eLFs contained only very few MOG_tet_^+^ B cells (Figure S6C, left panel). However, occasionally some eLFs displayed a massive accumulation of MOG_tet_^+^ B cells (Figure S6C, right panel). Together, these data indicate that transferred MOG-specific Th17 cells recruit B cells with the same antigen specificity from the endogenous pool and that maturation and/or expansion of MOG-specific B cells may at least partially happen in meningeal eLFs.

### CNS-infiltrating B cells respond to T cell cytokines

Our transcriptomic analyses suggest that many of our CNS B cells in Th17 adoptive transfer EAE are highly activated, poised for entering GC reactions and undergoing clonal expansion (Fig. 2 and 3). In addition to cellular interactions, phenotype and function can also be shaped by cytokines. Thus, we evaluated whether we could find evidence of cytokine-induced signaling in CNS B cells. B cells are known to strongly respond to cues from both type I and type II interferons and thus we compared gene expression in our clusters to publicly available gene signatures induced by IFNγ and IFNα. Although, some IFNγ is produced by transferred Th17 cells as well as by endogenous T cells in our model, among the CNS-specific clusters only cluster 6 and 7 showed evidence of IFNγ-driven signaling compared to the other CNS B cells, and this was enhanced in CNS B cells compared to peripheral B cells within the same cluster (Figure 4A). In addition, cluster 5 B cells, where only 32 % of cells are derived from CNS, express IFNγ-driven genes including *Stat1* and *Irf7* (Fig. S4A and Table S1). In contrast, the CNS-specific clusters did not show an enhanced response to IFNα neither compared to other CNS B cells nor compared to peripheral B cells within the same cluster (Figure S7A), suggesting that IFNα is not a relevant component of the CNS-specific cytokine milieu in our model. Among the CNS-enriched B cell clusters, cluster 7 and 9 exhibited elevated expression of genes induced by IL-21, and for all CNS-specific clusters expression was always higher in CNS B cells compared to peripheral B cells from the same cluster (Fig. 4B). Since IL-21 is produced by Th17 cells and is also a T helper cell-derived cytokine important for B cell responses in GC reactions, this speaks for T cells providing help to B cells to initiate GC reactions in the CNS. Indeed, expression levels of IL-21 induced genes were highest in cluster 7 and 9 (Figure 4B), which were shown to express markers of LZ and DZ GC B cells, respectively (Figure 2C, D). In line with the increased response to IL-21, flow cytometric analysis showed that CNS B cells also expressed higher levels of IL-21R than peripheral B cells (Figure 4C + S7C). CNS B cells also strongly responded to TNFα, while peripheral B cells did not (Figure 4D). This suggests that TNFα, which is produced by T cells but can also be released by myeloid cells and B cells themselves, is an important component of the CNS-specific cytokine milieu in our model. The highest levels of TNFα-induced gene expression were detected in cluster 7 B cells, which also showed very high levels of *Nr4a1* expression indicating that TNFα may even be a driver of B cell activation in CNS B cells. In line with this, we confirmed via flow cytometry that the frequency of activated CD69^+^ and CD83^+^ B cells was strongly increased in CNS B cells compared to splenic B cells (Figure 4E, F, S7B).

**Figure 4.**
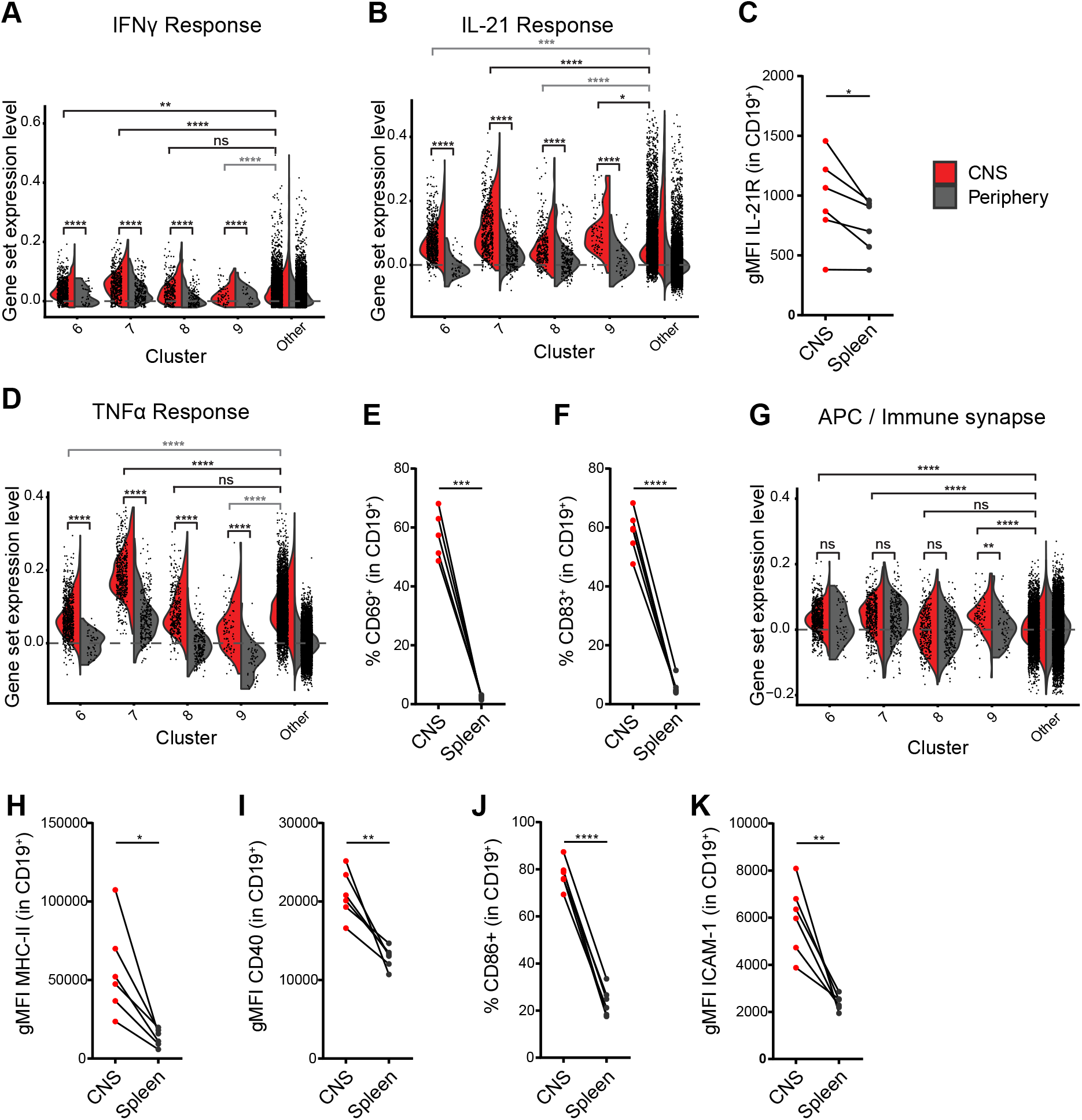
CNS-infiltrating B cells show elevated expression of genes associated with cytokine signaling, B cell activation and antigen presentation/immune synapse. (**A**, **B**, **D**, **G**) Violin plots showing expression levels of gene signatures induced by (**A**) IFNγ, (**B**) IL-21 and (**D**) TNFα, and (**G**) a gene set associated with antigen presentation/immune synapse. Two-sided *t*-test with FDR adjusted p-values. *p < 0.05; **p < 0.01; ***p < 0.001, ****p < 0.0001. Significantly downregulated gene sets are marked in grey. (**C**, **E**, **F**, **H-K**) At the peak of disease, cells were isolated from CNS or spleen and B cells were analyzed for expression of (**C**) IL-21R, (**E**) CD69, (**F**) CD83, (**H**) MHC-II, (**I**) CD40, (**J**) CD86 and (**K**) ICAM-1 (pre-gated on CD19^+^ cells) by flow cytometry. Graphs show the geometric mean fluorescence intensity (gMFI) of (**C**), IL-21R, (**H**) MHC-II, (**I**) CD40 or (**K**) ICAM-1 and the frequency of (**E**) CD69^+^, (**F**) CD83^+^ or (**J**) CD86^+^ cells. Paired *t*-test. *p < 0.05; **p < 0.01; ***p < 0.001, ****p < 0.0001. Dots represent individual mice.

### CNS B cells show enhanced APC function

Since we detected subpopulations of activated B cells that show features of GC B cells and respond to T cell cytokines, it is likely that eLF B cells closely interact with T cells in our model. To determine, whether CNS B cells not only respond to T cell help but also act on the T cells we determined the expression level of genes associated with APC function and immunological synapse (IS) formation in the CNS-specific clusters (Figure 4G). Notably, cluster 6, 7, and 9, which also showed the highest expression of the LZ and DZ GC B cell-associated genes, displayed highly significantly upregulated expression of APC/IS genes compared to other CNS B cells (Figure 4G). In line with this, CNS B cells showed highly elevated protein expression levels of MHC-II (Figure 4H, S7C), as well as elevated expression levels of the costimulatory molecule CD40 (Figure 4I, S7C), and increased frequencies of the costimulatory molecule CD86 (Figure 4J, S7B), which together with MHC-II form the central supramolecular activation complex (SMAC) of the immunological synapse. Furthermore, CNS B cells also showed increased levels of ICAM-1 (Figure 4K, S7C), which binds to LFA-1 on T cells and forms the peripheral SMAC of the IS. Taken together, our transcriptomic and flow cytometric analysis suggests that CNS B cells show enhanced capacity for antigen presentation, co-stimulation and formation of immunological synapses.

### T cells form long-lasting contacts with B cells in meningeal eLFs

Meningeal eLFs have been proposed to perpetuate chronic inflammation in the CNS compartment by providing a favorable environment for the interaction of T and B cells. This view is supported by our phenotypic analyses of CNS B cells that revealed their enhanced capacity for interaction with T cells. However, to our knowledge, it has never been demonstrated whether and to what effect T and B cells interact in meningeal eLFs. Therefore, we employed intravital two-photon microscopy to visualize the behavior of T and B cells in meningeal eLFs *in situ* and, in particular, to determine whether B cells are able to reactivate T cells (Figure 5A). To fluorescently label T cells, we used the ratiometric calcium sensor Twitch-2B, which allows for studying T cell activation in real-time by visualizing the increase of the intracellular calcium concentration upon TCR ligation (Thestrup et al., 2014). MOG-specific Th17 cells were transduced with a retrovirus containing Twitch-2B during differentiation, and we found that transduction efficiency was highest after restimulation with up to 60 % Twitch-2B-expressing Th17 cells (Figure S8A). Transduced Th17 cells were transferred into Mb1.Cre x ROSA26 tdTomato mice, which express tdTomato specifically in B cells (Figure 5A), and we confirmed that the transduced Th17 cells survived well *in vivo,* infiltrated the CNS, and showed stable expression of both Twitch-2B^+^ (Figure S8B) and effector cytokines in the CNS (Figure S8C). Furthermore, CNS-infiltrating B cells in Mb1.Cre x ROSA26 tdTomato Th17-EAE mice faithfully expressed tdTomato detected by flow cytometry as well as confocal microscopy of CNS cryosections (Figure S8D). In contrast, other CNS-infiltrating APCs such as CD11b^+^ macrophages showed frequencies of tdTomato^+^ cells below 4 % (not shown). In all diseased Th17 recipients, intravital microscopy of spinal cord showed massive accumulation of Twitch-2B^+^ T cells and clustered tdTomato^+^ B cells in the meninges, revealing eLFs of substantial size and structure (Figure 5B). Since in this experiment, the transduction rate of T cells ranked at 58 %, it can be assumed that the accumulation of transferred T cells is even underestimated in this image. Zooming inside eLFs revealed extensive T:B cell contacts within these structures (Figure 5C). To compare the T cell response of the MOG-specific 2D2 Th17 cells to non-encephalitogenic Th17 cells, a set of control experiments was performed, where ovalbumin (OVA)-specific OT-II Th17 cells were transduced with the Twitch-2B sensor and transferred together with non-transduced 2D2 Th17 cells into Mb1.Cre x ROSA26 tdTomato mice. Both 2D2 and OT-II T cells were investigated regarding their interaction with B cells by monitoring the movement of individual cells in the 3D follicle structure over time. Intriguingly, 2D2 T cells interacted more intensely with B cells than OT-II T cells. Around half of the 2D2 T cells formed contacts with B cells compared to only 15 % of OT-II T cells (Figure 5D). 2D2 T cells and B cells engaged in very long contacts, which often lasted during an entire observation period of 35 min. In contrast, OT-II T cells formed B cell contacts of much shorter duration (Figure 5E). Setting a threshold of five minutes to define short and long contacts, as was suggested by measuring the contact length of B cells and Tfh cells in regular GC reactions (Jacobsen et al., 2021), we found that the majority of 2D2 T cells (82 %) engaged in long T:B cell contacts compared to only a third of OT-II T cells (Figure 5F). Accordingly, 2D2 T cells engaged in T:B cell contacts also showed a reduced velocity (mean 1.63 µm/min) compared to 2D2 T cells without B cell contact (mean 2.47 µm/min), while OT-II T cells exhibited significantly higher motility (mean 3.00 µm/min) than both 2D2 T cell groups (Figure 5G). Taken together, this indicates that MOG-specific 2D2 T cells are being arrested due to cognate interaction with B cells.

**Figure 5.**
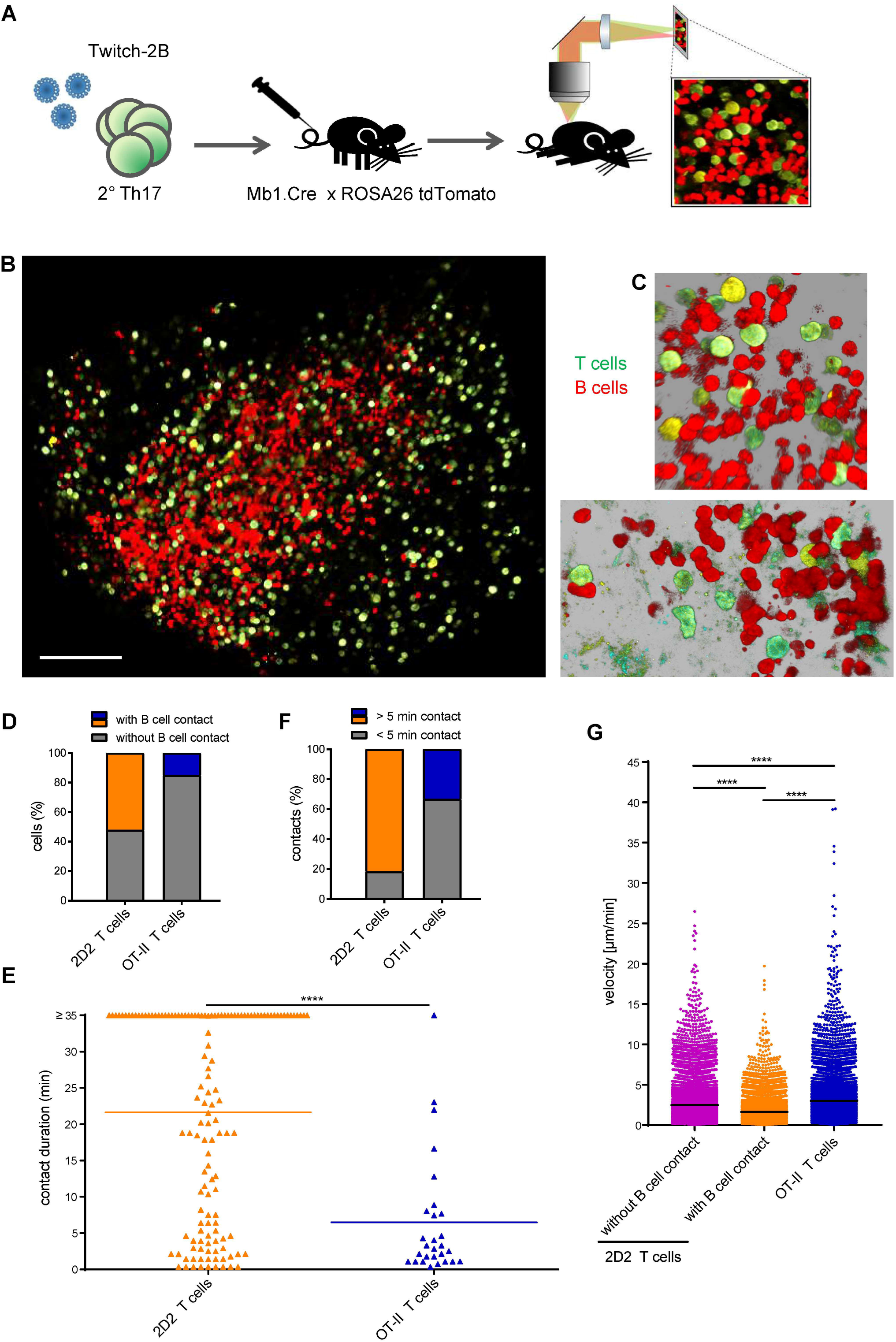
2D2 T cells form long-lasting contacts with B cells compared to OT-II T cells. Intravital microscopy of Twitch-2B^+^ T cells and tdTomato^+^ B cells in spinal cord eLFs of Th17-EAE mice. (**A**) Experimental outline. MOG-specific T cells from 2D2 mice are retrovirally transduced with the Twitch-2B construct during their differentiation into Th17 cells. Upon adoptive transfer in Mb1.Cre x ROSA26 tdTomato mice, intravital two-photon microscopy of spinal cord eLFs in EAE mice can be performed to visualize interactions of Th17 cells (green) and B cells (red), as well as subsequent activation events of Th17 cells. (**B**) Dorsal view onto eLF containing Twitch-2B^+^ T cells (green) and tdTomato^+^ B cells (red). Scale bar: 100µm. (**C**) Exemplary images of T:B cell contacts. (**D**) Proportion of 2D2 (n = 232 cells from six independent experiments) and OT-II T cells (n = 107 cells from four independent experiments) forming contact with B cells. (**E**) Quantification of T:B cell contact duration. For both experimental groups, videos of 15-50 min length were acquired. Dots represent individual contacts. Horizontal lines indicate means. Mann-Whitney U test. ****p < 0.0001. (**F**) Proportion of long (> 5 min) and short (< 5 min) T:B cell contacts. (**G**) Quantification of the velocity of 2D2 T cells without B cell contact (n = 10938 velocities), 2D2 T cells with B cell contact (n = 8314 velocities) and OT-II T cells (n = 9413 velocities). One-way ANOVA with Tukey post-test. ****p < 0.0001. Horizontal lines indicate means. Graph shows cumulative data from (2D2) six or (OT-II) four independent experiments.

### B cells reactivate T cells in meningeal eLFs

To investigate whether 2D2 T cells become activated during B cell contact, we made use of the ratiometric calcium sensor Twitch-2B, which represents a powerful tool to visualize intracellular calcium signaling in activated T cells. Importantly, many 2D2 T cells showed elevated calcium levels and, in addition, appeared very blastic, indicating that they are highly activated (Figure 6A, Movie 1). Quantitative analysis of the calcium ratios revealed that 2D2 Th17 cells exhibited highly increased calcium levels compared to OT-II Th17 cells. To correctly interpret increased intracellular calcium concentrations as calcium signaling, we set a threshold at the maximal level reached spontaneously by 95 % of OT-II T cells. For 2D2 T cells, 75.0 % of calcium ratios in T cells with B cell contact exceeded the threshold compared to 47.2 % of calcium ratios in T cells without B cell contact, showing that T cells exhibit more pronounced calcium signaling during T:B cell contact (Figure 6B). Notably, the majority of 2D2 T cells (81.7 %) exhibited calcium spikes compared to less than a third of OT-II T cells (Figure 6C). 2D2 T cells displayed long-lasting calcium signaling, which could often be observed during an entire observation period of 35 min, whereas OT-II T cells primarily showed calcium signaling of short duration (Figure S8E). To define short- and longterm calcium spikes, we used a cut-off at two minutes since calcium spikes longer than two minutes require antigen-dependent T cell activation (Kyratsous et al., 2017). We found that about 60% of 2D2 T cells showed long-term signaling compared to less than 7 % of OT-II T cells. 2D2 T cells also showed more short-term signaling than OT-II cells, whereas the majority of OT-II cells showed no calcium signaling at all (Figure 6D). The difference in T cell response upon B cell contact became even more evident by comparing single cell tracks of 2D2 and OT-II T cells. 2D2 T cells exhibited pronounced calcium signaling during B cell contact (Figure 6E, S8F), whereas calcium ratios of OT-II T cells remained unchanged during B cell contact (Figure 6F, S8G). For better visualization of T:B cell interaction and the ensuing T cell response, Figure 6G depicts snapshots of representative 2D2 and OT-II T cells during B cell contact. The first 2D2 T cell formed a long-lasting contact with one and later two B cells. While the cell appeared to be arrested and only underwent minor changes in cell shape, it displayed high calcium signaling for over 30 min. In the second example, the 2D2 T cell showed low calcium levels before contacting the B cell and increasing calcium levels upon contact formation. In contrast, the OT-II T cell did not show a difference in calcium levels before and during contact with a B cell (1.4 min - 14.8 min).Together, these data indicate that B cells in eLFs provide a reactivation stimulus to 2D2 T cells, but not to OT-II T cells.

**Figure 6.**
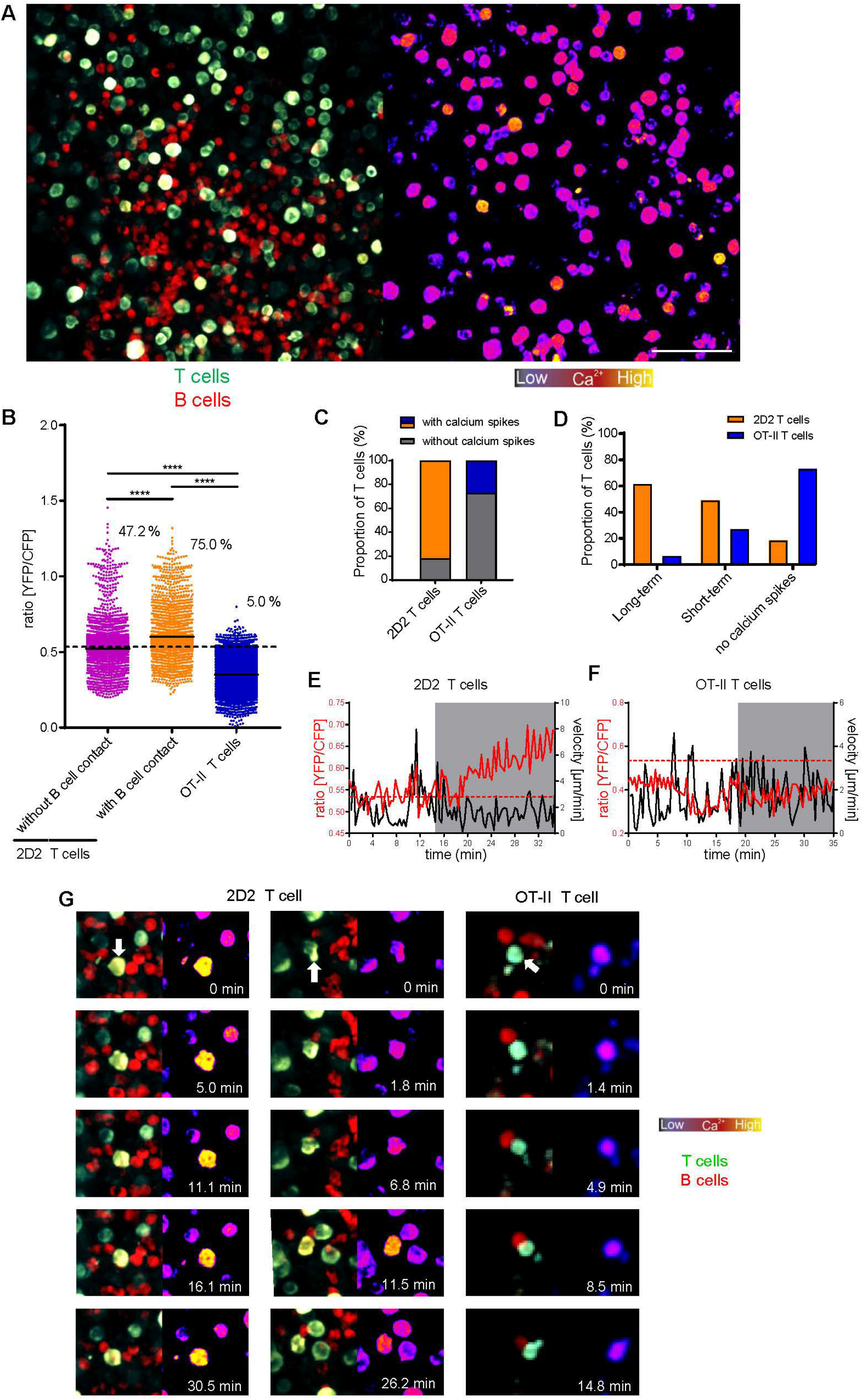
2D2 T cells, but not OT-II T cells, become reactivated during B cell contact. Intravital microscopy of Twitch-2B^+^ T cells and tdTomato^+^ B cells in spinal cord eLFs of Th17-EAE mice. (**A**) Single time point of a representative video. (Left) Fluorescence overlay of Twitch-2B^+^ T cells (green) and tdTomato^+^ B cells (red) and (right) pseudocolor calcium ratio image showing calcium levels of T cells. Scale bar: 50 µm. (**B**) Quantification of the calcium indicator ratio (YFP/CFP) of 2D2 T cells without B cell contact (n = 4988 ratios), 2D2 T cells with B cell contact (n = 3653 ratios) and OT-II T cells (n = 9531 ratios). One-way ANOVA with Tukey post-test. ****p < 0.0001. The black dashed line indicates the ratio threshold (0.534). Horizontal lines indicate means. The graph shows cumulative data from (2D2) three or (OT-II) four independent experiments. (**C**) Proportion of 2D2 and OT-II T cells exhibiting calcium signaling. (**D**) Proportion of T cells with short-term (< 2 min), long-term (> 2 min) or no calcium spikes. (**C**, **D**) Cumulative data of 2D2 (n = 104 cells from three independent experiments) and OT-II T cells (n = 107 cells from four independent experiments). (**E, F**) Representative track of (**E**) 2D2 T cells (n = 104 cells from three independent experiments) and (**F**) OT-II T cells (n = 107 cells from four independent experiments) showing calcium indicator ratio (YFP/CFP, red) and velocity (black) in each time frame. The red dashed line indicates the ratio threshold (0.534). Grey shading highlights time frames with B cell contact. (**G**) Exemplary microscopic images of (left and middle panel) two representative 2D2 and (right panel) a representative OT-II T cell. (Left) Fluorescence overlay of Twitch-2B^+^ T cells (green) and tdTomato^+^ B cells (red) and (right) pseudocolor calcium ratio image showing calcium levels of T cells. White arrow indicates cell of interest. (Left panel) 2D2 T cell has B cell contact at all indicated time points. (Middle panel) 0 min: before B cell contact; 1.8 min – 26.2 min: with B cell contact. (Right panel) 0 min: before B cell contact; 1.4 min – 14.8 min: with B cell contact.

### T cells depend on CNS B cells to sustain pro-inflammatory effector functions in the CNS

Intravital imaging showed long-lasting T:B cell contacts in eLFs that lead to increased Ca^2+^ signaling indicative of T cell reactivation. To investigate the functional consequences of these interactions, we transferred 2D2 Th17 cells into B cell-deficient Mb1-KO mice. At the peak of disease, infiltrating cells were recovered from the CNS, and T cells were analyzed for their cytokine profile. Transferred T cells could be distinguished from endogenous T cells due to their expression of the transgenic MOG-specific Vα3.2 TCR chain. Interestingly, the frequency of IL-17A^+^, IFNγ^+^ and IL-17A^+^IFNγ^+^ producers was significantly decreased among transferred T cells in Mb1-KO mice compared to WT mice. Similarly, endogenous T cells, which mainly produced IFNγ, showed lower levels of IL-17A^+^, IFNγ^+^ and IL-17A^+^IFNγ^+^ cells in Mb1-KO mice (Figure 7A). However, in the spleen we did not find decreased T cell effector cytokines in the absence of B cells (Figure 7B), strengthening the importance of the reactivation stimulus provided by B cells in the eLFs in the CNS rather than in the periphery.

**Figure 7.**
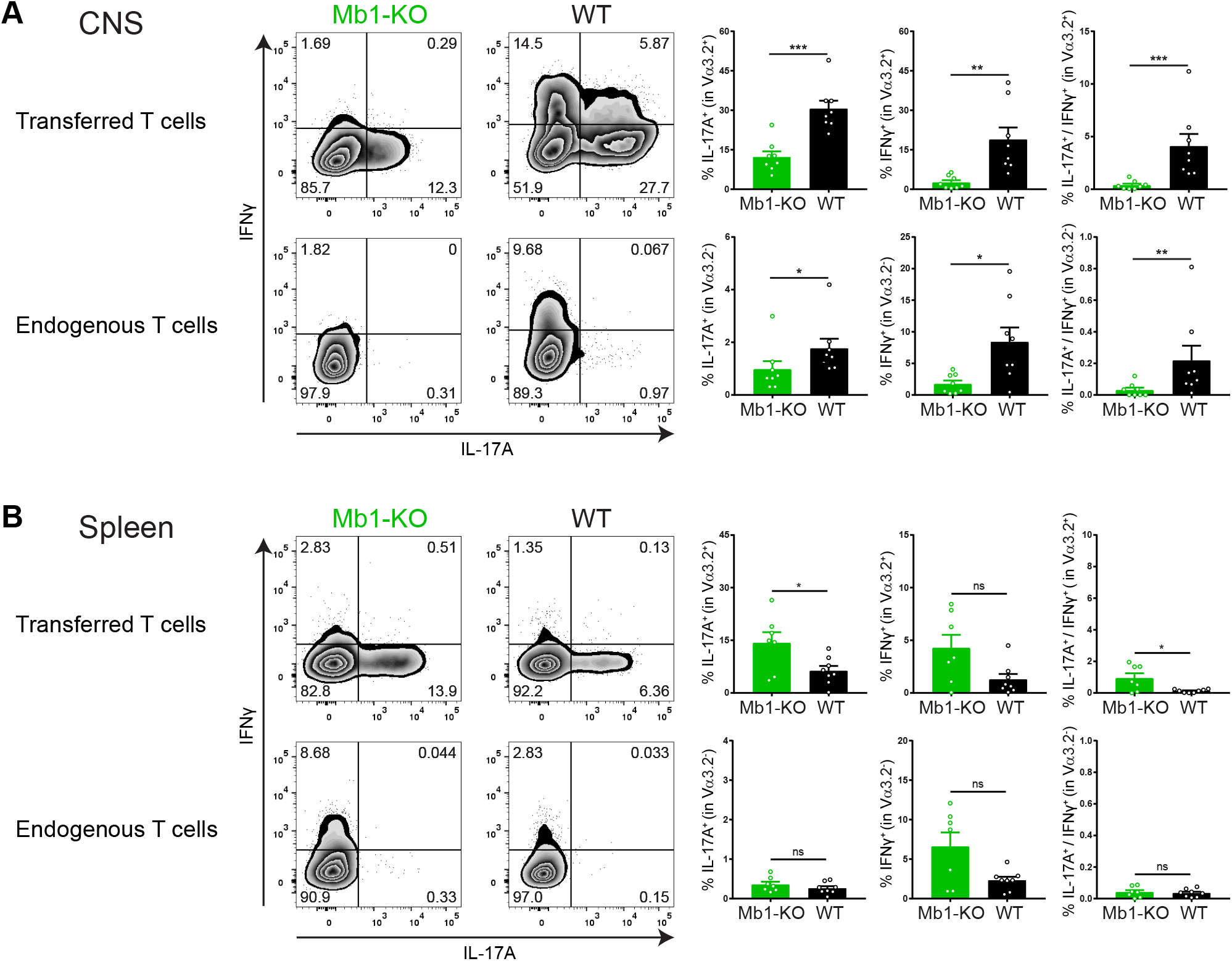
T cells depend on B cells to maintain a highly pro-inflammatory cytokine profile in the CNS. Comparison of T cell responses in Th17-mediated EAE in B cell-deficient Mb1-KO mice and WT mice. At the peak of disease, cells were isolated from (**A**) CNS or (**B**) spleen and T cells were analyzed for IL-17A and IFNγ expression by intracellular cytokine staining. Transferred T cells were identified by their expression of the transgenic MOG-specific Vα3.2 TCR. Representative flow cytometry plots and quantification of IL-17A^+^, IFNγ^+^ and IL-17A^+^IFNγ^+^ producers of (upper panel) transferred (Vα3.2^+^) and (lower panel) endogenous (Vα3.2^-^) T cells. Graphs show cumulative data from two independent experiments. (**A**, upper panel) unpaired *t*-test (left), Welch’s t-test (middle), Mann-Whitney U test (right). (**A**, lower panel) Mann-Whitney U test (left and right), Welch’s t-test (middle). (**B**, upper panel) unpaired *t*-test (left), Mann-Whitney U test (middle), Welch’s t-test (right). (**B**, lower panel) unpaired *t*-test (left and right), Welch’s t-test (middle). *p < 0.05; **p < 0.01; ***p < 0.001. Mean ± SEM. Dots represent individual mice.

## Discussion

In our study, we combined flow cytometric, transcriptomic and imaging analyses to investigate the unique relationship of Th17 cells and B cells and, in particular, to shed light on their cooperation in meningeal eLFs. In our Th17 adoptive transfer EAE model, which features formation of large eLFs within the meninges of brain and spinal cord, these follicles contain clusters of proliferating T and B cells, show expression of germinal center markers, and are reminiscent of eLFs in MS patients. Through single-cell RNA sequencing and flow cytometric analyses we discovered that substantial subsets of CNS-infiltrating B cells are highly activated, poised for entering GC reactions and undergoing clonal expansion. In addition, CNS B cells respond to T cell cytokines and show enhanced capacity for antigen presentation, co-stimulation and formation of immunological synapses. Correspondingly, intravital imaging provided the first real-time observation of intense T:B cell cooperation in meningeal eLFs and demonstrated that T and B cells form antigen-dependent long-lived contacts, whereby B cells provide a reactivation stimulus to T cells. Consequently, we showed that this reactivation stimulus from B cells is required for CNS T cells to maintain their highly pro-inflammatory cytokine profile.

Histological analyses in the Th17 adoptive transfer EAE model suggest that eLFs appear in different stages of maturation, even within the same animal, and that many eLFs are not fully matured at the time point of investigation (Figure 1). Single-cell transcriptomic analysis of CNS B cells largely confirmed this observation, as we identified several clusters of substantial size of B cells expressing markers of activation and markers associated with light zone GC B cells, but only very few dark zone GC B cells (Figure 2). Therefore, we suggest that our model features mostly eLFs harboring very early stages of GC reactions and this is probably due to the rather short disease duration in our model: since B cells only start accumulating in the CNS at the beginning of clinical symptoms, they had only 4-7 days to arrive and mature in eLFs until the recipients were analyzed. We speculate that the GC reaction in CNS eLFs would progress to more advanced stages with time, however, due to the severe disease course these later stages cannot be studied in our model. Nonetheless, we found elevated titers of MOG-specific antibodies in the serum at the late peak of disease compared to very low levels before disease onset (Figure S6A) and some eLFs containing massive amounts of MOG-specific B cells (Figure S6C). This shows the capacity of eLFs to support initial priming and maturation or -at the very least - expansion of autoreactive B cells that may have been primed in the periphery and then migrated to eLFs to expand. This speaks for an active disease-promoting role of eLFs since CNS-reactive autoantibodies are known to be pathogenic in CNS autoimmunity (Mader et al., 2020). In addition, eLFs have been suspected to mediate cortical damage in the underlying parenchyma by releasing soluble pro-inflammatory factors, which may include CNS-reactive antibodies but also cytokines. In fact, we found CNS B cells to respond strongly to TNFα (Figure 4D) and IL-21 (Figure 4B), an important Th17 effector cytokine, suggesting that the eLF cytokine milieu is enriched for those cytokines in comparison to the periphery. Especially TNFα has pleiotropic effects on microglia, neurons, astrocytes and oligodendrocytes (Raffaele et al., 2020), is increased in the CSF of MS patients with high levels of meningeal inflammation and grey matter damage (Magliozzi et al., 2018), and is therefore a prime candidate for further investigation regarding the association of eLFs and cortical damage. Although IL-21 is primarily known to act on immune cells, IL-21 receptor is also expressed by cortical neurons and expression is up-regulated in MS patients (Tzartos et al., 2011). In a stroke model, IL-21 signaling in neurons was associated with increased autophagy (Clarkson et al., 2014), suggesting that IL-21 may also be an interesting candidate for mediating cortical damage underneath eLFs. The effect of other Th17-derived cytokines on B cells such as IL-17A was not investigated, since there are no gene expression signatures available. Surprisingly, IFNγ- driven signaling seemed less important for CNS B cells (Figure 4A), although some IFNy is produced by transferred Th17 cells, as well as by host-derived endogenous T cells in the CNS (Figure S1D). However, the effect of IFNγ on CNS B cells may be more pronounced in case of Th1 adoptive transfer EAE, where the CNS T cells secrete massive amounts of IFNγ (Figure S1C) but much fewer eLFs are formed.

Notably, our study revealed the presence of B1a / MZ-like B cells in the CNS (Figure 2E-H, S5D-F), which may result from migration of B1a / MZ B cells from the spleen or peritoneum to the CNS. However, it is also possible that the inflamed CNS provides a microenvironment, which induces infiltrating B cells to adopt a B1a / MZ-like phenotype. In several autoimmune settings, MZ-like B cells infiltrate the target tissue, e.g. the thyroid gland (Segundo et al., 2001), or the salivary glands (Groom et al., 2002). Interestingly, a MZ-like population was also detected in CSF from MS patients suggesting that a human counterpart of the cluster 8 B1a / MZ-like B cells may also exist in MS patients (Lee-Chang et al., 2011). Our B1a / MZ-like B cells expand clonally (Figure 3A, C), but show less evidence of a T cell help-induced GC reaction (Figure 2C, D), which is in line with the property of regular B1a / MZ B cells to operate more independently of T cells. Considering the high expansion factors and expression levels of plasmablast-associated genes (Figure 3C, D), they are probably a rich source of antibodies, but it remains to be determined whether these are autoreactive. It is important to note that normal B1a and MZ B cells can also perform regulatory functions (Zhang, 2013). They are more potent IL-10 producers compared to other splenic B cell subsets (Barr et al., 2012) and may play a protective role in autoimmunity (Lenert et al., 2005; Matsushita et al., 2018). In fact, several upregulated genes in cluster 8, such as *Fcrl5*, *Zbtb20*, *Cd9* and *Apoe* (Table S1), were recently defined as markers of regulatory B cells by single-cell RNA sequencing of B cells from different murine organs (Yang et al., 2021). Therefore, it is possible that they play a regulatory function in the CNS, especially in later disease stages and when differentiating into plasma cells.

The importance of B cells as APCs in neuroinflammation has been reported in several EAE studies (Bettelli et al., 2006; Krishnamoorthy et al., 2006; Molnarfi et al., 2013; Parker Harp et al., 2015; Weber et al., 2010). In our model, we demonstrated that CNS B cells reactivate MOG-specific T cells in eLFs, as shown by intravital microscopy (Figure 6B, E, G) and adoptive transfer into B cell-deficient Mb1-KO mice (Figure 7). However, considering that the B cell repertoire in our WT recipient mice is polyclonal and only few eLFs were positive for MOG-specific B cells by MOG_tet_-staining (Figure S6), B cells in eLFs cannot all be MOG-specific. Thus, we speculate that antigen bound to MHC-II is transferred to B cells from other APCs, which are also present in eLFs (Figure S2). Although, the mechanism requires further investigations, this could happen for example by trogocytosis (Schriek et al., 2022), or via extracellular vesicles (Becker et al., 2021) derived from rare MOG-specific B cells or from ubiquitous APCs including macrophages and DCs. Due to the extensive tissue damage at this late peak of disease the area must be saturated with released CNS autoantigens ready to be taken up and interestingly, extracellular vesicles containing MOG were also shown to be increased in the CSF of MS patients during disease activity (Galazka et al., 2018). Another point of consideration is that the 2D2 TCR does not exclusively recognize MOG, as it was shown to also respond to neurofilament (Krishnamoorthy et al., 2009). Nonetheless, we demonstrated that OT-II T cells do not become reactivated by B cells in eLFs even though they are also differentiated into Th17 cells (Figure 6B-D, F, G), indicating that the reactivation stimulus provided by the B cells to the 2D2 T cells must be CNS antigen-dependent.

The clinical significance of eLFs requires further investigation. In MS, formation of eLFs is associated with a more severe disease course and subpial cortical as well as spinal cord pathology (Howell *et al*., 2011; Magliozzi *et al*., 2007; Magliozzi *et al*., 2010; Reali *et al*., 2020; Serafini *et al*., 2007; Serafini *et al*., 2004). Due to the short and severe disease course, our Th17-transfer EAE model is not suitable to study the clinical impact of eLFs, but in the spontaneous opticospinal EAE model, which features both genetically modified MOG-specific T and B cells and a milder disease course that allows for longer observation periods, a correlation between eLF presence and a more severe disease course was found (Dang *et al*., 2015). In the same model, a recent study suggested an immunoregulatory function of eLFs since eLF abrogation resulted in a higher disease burden (Mitsdorffer et al., 2021). It is possible that eLFs predominantly form in severe disease cases in an attempt to contain excessive inflammation, and this would involve presence of regulatory T_FR_ cells and/or regulatory B cells producing anti-inflammatory cytokines. Although, Foxp3^+^ regulatory T_FR_ cells were not detected in eLFs in post-mortem CNS tissue from MS patients (Bell et al., 2019), Mitsdörffer and colleagues identified B cells producing the immunoregulatory cytokines IL-10 and IL-35 in eLFs in the opticospinal EAE model (Mitsdorffer *et al*., 2021). Although we did not detect IL-10 or IL-35 in our transcriptomic dataset, we also found markers of regulatory B cells expressed in the B1a / MZ-like cluster 8 B cells. Thus, eLFs may provide a supportive environment for regulatory B cells and this may become more relevant at later time points, as differentiation to plasma cells, which are a known source of anti-inflammatory cytokines, progresses further. eLFs with disease-promoting and regulatory impact may co-exist in MS either in different subsets of patients or in different phases of the disease. Therefore, both models are important to study the role of eLFs in CNS autoimmunity, as each of them may represent one end of the spectrum of what can happen within eLFs, ranging from immunoregulation to immunopathology.

In contrast to the opticospinal EAE model, Th17-mediated EAE employs non-transgenic B cells with an endogenous repertoire and therefore, the B cell response in our model can be assumed to better reflect B cell-related disease aspects in MS. The possibility that eLFs support initial priming and maturation and/or expansion of CNS-reactive B cell clones has clear immunopathological implications and the increased MOG-specific antibody titers and rare MOG_tet_-positive eLFs in our model point in that direction, although more research is required to confirm eLFs as sources of CNS-reactive B cells. In addition, our data suggests that -zmaybe even independent of their own specificity - B cells in eLFs excel at presenting CNS antigen to interacting T cells and reactivating them to maintain pro-inflammatory effector functions. Taken together, we conclude that the bilateral interactions of T and B cells in meningeal eLFs in our model support smoldering inflammatory processes that could drive CNS pathology even behind a closed blood brain barrier, and propose that further research investigating whether T:B cell cooperation in eLFs can be therapeutically targeted is warranted.

## Materials and methods

### Mice

C57BL/6 mice were purchased from Janvier Labs. C57BL/6 mice with a transgenic TCR specific for the MOG_35-55_ peptide, here referred to as 2D2 mice, have been described previously (Bettelli et al., 2003) and were purchased from The Jackson Laboratory. IgH^MOG^ mice on a C57BL/6 background, also known as Th mice, carrying an Ig heavy chain knock-in of the rearranged VDJ gene from the MOG-specific monoclonal antibody 8.18C5, have been described previously (Litzenburger et al., 1998) and were kindly provided by Prof. Hartmut Wekerle. Mb1-Cre mice on a C57BL/6 background carry a codon optimized Cre recombinase, replacing exon 2 and 3 of the *Cd79a* gene, which encodes the Ig-α subunit of the BCR and is exclusively expressed in B cells. This knock-in leads to abolishment of endogenous *Cd79a* gene function and expression of Cre recombinase under control of the *Cd79a* promoter (Hobeika et al., 2006). Mb1-Cre mice were purchased from The Jackson Laboratory. Homozygous mice are B cell-deficient and hereafter referred to as Mb1-KO mice. Mb1-Cre mice were crossed to ROSA26 tdTomato mice to obtain mice that are heterozygous for both Mb1-Cre and ROSA26 tdTomato, thereby expressing tdTomato specifically in B cells for intravital microscopy experiments. ROSA26 tdTomato mice on a C57BL/6 background carry a *loxP*-flanked STOP cassette and *tdTomat*o gene, both inserted into the ROSA26 locus. Upon Cre-mediated recombination, tdTomato fluorescence is robustly expressed in these mice (Madisen et al., 2010). ROSA26 tdTomato mice were purchased from The Jackson Laboratory. OT-II mice, carrying a transgenic TCR specific for the ovalbumin_323-339_ peptide on a C57BL/6 background (Barnden et al., 1998), were purchased from The Jackson Laboratory.

All animals used in this study were housed and bred in specific pathogen-free conditions in the Core Facility Animal Models at the Biomedical Center of the Ludwig-Maximilians-Universität München. Animal experiments were designed and performed in accordance with regulations of the animal welfare acts and under approval by the animal ethics committee of the state of Bavaria (Regierung von Oberbayern) in accordance with European guidelines.

### Isolation of cells from spleen and lymph nodes

Spleens were mashed and centrifuged at 300 g for 5 minutes (min) at 4 °C. To lyse erythrocytes, the cells were resuspended in ACK buffer (150 mM NH4Cl (Merck), 10 mM KHCO3 (Merck), 0.1 mM EDTA (Sigma)) and incubated for 1.5 min at RT. The reaction was stopped by adding 9 ml T cell medium (RPMI 1640 (Sigma) supplemented with 10 % heat-inactivated FBS, 1 % penicillin-streptomycin, 10 mM HEPES, 2 mM L-glutamine, 1 % non-essential amino acids, 1 mM sodium pyruvate and 50 µM β-mercaptoethanol), and the cell suspension was passed through a 70 µm cell strainer. After centrifugation, the cells were resuspended in T cell medium. Lymph nodes (LNs) were cut into pieces, mashed and filtered through a 70 µm cell strainer. Then, the cells were centrifuged at 300 g for 5 min at 4 °C and resuspended in T cell medium.

### *In vitro* differentiation of T effector cells

Naïve CD4^+^ T cells were purified from spleen and lymph nodes of 6-15 week old 2D2 mice or OT-II mice using the naïve CD4^+^ T cell isolation Kit (Miltenyi Biotec) according to the manufacturer’s instructions. After isolation of naïve CD4^+^ T cells the rest of the cells (flow-through from T cell purification) was irradiated at 35 Gy and used as APCs for costimulation of differentiating T cells. Naïve T cells were cultured at a concentration of 1.5-2×10^6^ ml^-1^ in T cell medium in the presence of 7.5-10×10^6^ ml^-1^ irradiated APCs (ratio T cells : APCs is 1 : 5) and 2.5 μg/ml soluble anti-CD3 antibody (clone 145-2C11, BioXCell). Th1 cells were generated by addition of IL-12 at a concentration of 10 ng/ml and anti-IL-4 antibody (clone 11B11, BioXCell) at a concentration of 10 µg/ml into the culture. For the generation of Th17 cells, naïve T cells were cultured with IL-6 at a concentration of 30 ng/ml, TGFβ at a concentration of 3 ng/ml, IL-1β at a concentration of 20 ng/ml, and anti-IFNγ (clone XMG1.2, BioXCell) and anti-IL-4 antibody (clone 11B11, BioXCell) at a concentration of 10 µg/ml. After 48h, Th1 cells and Th17 cells were split with medium containing 10 ng/ml of IL-2 and medium containing 10 ng/ml of IL-23, respectively. All cytokines were purchased from Biolegend except IL-23 (Miltenyi Biotec). The different T cell subsets were analyzed for cytokine production after 4 days by intracellular cytokine staining and subsequent flow cytometry. After 5-8 days of primary culture, T cells reached a resting state and were restimulated at a concentration of 2×10^6^ ml^-1^ for 48 h in the presence of plate-bound anti-CD3 (clone 145-2C11, BioXCell) and anti-CD28 (clone PV-1; BioXCell) antibodies both at 2 µg/ml in fresh medium without any cytokines.

### Retrovirus production

Phoenix-eco cells (Pear et al., 1993) were routinely cultured in TCM medium (DMEM (Sigma) containing 10 % heat-inactivated FBS, 1 % penicillin-streptomycin, 2 mM L-glutamine, 360 mg/l asparagine, 1 % non-essential amino acids, 1 mM sodium pyruvate and 57.2 µM β-mercaptoethanol). The cells were kept in a subconfluent state by splitting them every 2-3 days. For production of retroviral particles, the cells were plated at 1.5-2×10^6^ per 10 cm dish in 10 ml TCM medium and allowed to attach for 18-24 hours. First, chloroquine diphosphate (Sigma) was added at a concentration of 25 µM. Per dish, 12 µg pMSCV-Δneo-Twitch-2B plasmid (kindly provided by Dr. Naoto Kawakami) and 3.5 µg pCL-Eco packaging plasmid (kindly provided by Dr. Gurumoorthy Krishnamoorthy) were prepared in 438 µl H_2_O. Subsequently, 62 µl 2 M CaCl_2_ (Riedel de Haën) were added followed by dropwise addition of 500 µl 2X BES (50 mM N,N-bis(2hydroxyethyl)-2-aminoethanesulfonic acid (Sigma) + 280 mM NaCl (Sigma) + 1.5 mM Na_2_HPO_4_ (Roth) in H_2_O) while vortexing. This mixture was incubated for 20 min at 37 °C and then added dropwise to the cells. On the next day, the medium was replaced with 8 ml fresh and warm TCM medium. After another 24 hours, the cell culture supernatant was collected using a 10 ml syringe, filtered through a 0.45 µm filter and kept at 4 °C. To every dish, 4 ml fresh and warm TCM medium was added and the cells were incubated for another day. The supernatant was collected again, filtered through a 0.45 µm filter and pooled with the supernatant from the day before. Then, the supernatant was concentrated by centrifugation at 1000 g at 4 °C through Amicon® Ultra 15 mL Centrifugal Filters (100 K; Merck). Finally, the concentrated supernatant was snap frozen and stored at −80 °C.

### Retroviral transduction of Th17 cells

#### Retroviral transduction during primary phase

After 48 h of primary culture, Th17 cells were split (usually 1:3) and transduced by adding 1:80-diluted retroviral supernatant and 4 µg/ml polybrene (Sigma) in addition to 10 ng/ml IL-23 and 10 µg/ml anti-IFNγ antibody (all diluted in pre-warmed T cell medium and calculated for culture volume). Then, cells were spin-infected at 480 g for 90 min at 30 °C. On the next day, half of the medium was replaced with fresh and warm T cell medium containing 10 ng/ml IL-23 and 10 µg/ml anti-IFNγ antibody. 48 h after transduction, Twitch-2B expression was analyzed via GFP-signal by flow cytometry.

#### Retroviral transduction during secondary phase

24 h after restimulation with anti-CD3/CD28 antibodies, half of the medium was taken up and Th17 cells were transduced by adding 1:80-diluted retroviral supernatant and 4 µg/ml polybrene (Sigma) (all diluted in pre-warmed T cell medium and calculated for culture volume). Then, cells were spin-infected at 480 g for 90 min at 30 °C. On the next day, half of the medium was replaced with fresh and warm T cell medium. If cells were used for EAE experiments, cells were collected and washed for adoptive transfer, while an aliquot of cells, with half of the medium being replaced with fresh and warm T cell medium, was kept for determining the transduction efficacy. 48 h after transduction, Twitch-2B expression was analyzed via GFP-signal by flow cytometry.

### Adoptive transfer EAE

Naïve MOG-specific CD4^+^ T cells, isolated from spleen and lymph nodes of 2D2 mice, were differentiated into Th1 or Th17 cells and restimulated with anti-CD3 and anti-CD28 antibodies. 48 h after restimulation, cells were collected and washed three times with PBS. In Th1-EAE, 1.7-3.2×10^6^ IFNγ-producing cells were injected intraperitoneally (i.p.) into C57BL/6 recipients. In Th17-EAE, 4×10^6^ IL-17A-producing cells were injected i.p. or intravenously (i.v.) into C57BL/6 or Mb1-KO or Mb1Cre x ROSA26 tdTomato recipients. In case of 2D2 and OT-II T cell co-transfer, 3×10^6^ IL-17A-producing 2D2 cells and 2×10^6^ IL-17A-producing OT-II cells were injected together i.p. into Mb1Cre x ROSA26 tdTomato recipients. Animals of both genders were used as donors, whereby male cells were injected only into male recipients. Clinical disease developed 8-15 days post transfer. Mice were monitored daily for the development of clinical signs of EAE, including classical and atypical disease signs, according to the following criteria: 0, no disease; 0.5, decreased tail tonus; 1, limp tail or mild balance defects; 1.5, limp tail and ataxia; 2, hind limb weakness, or severe balance defects that cause spontaneous falling over; 2.5, partial hind limb paralysis; 3, complete hind limb paralysis or very severe balance defects that prevent walking; 3.5, complete hind limb paralysis and partial front limb paralysis; 4, complete front and hind limb paralysis or inability to move body weight into a different position; 5, moribund state.

### Isolation of infiltrating mononuclear cells from the CNS

Recipient mice, which had been on a high score for several days to allow for continued immune cell infiltration, were sacrificed and perfused through the left cardiac ventricle with PBS. Brains were dissected by removing the skull using forceps and scissors, while spinal cords were flushed out from the bone column using a 21 gauge needle attached to a syringe filled with PBS. In addition, the vertebral column was opened to collect parts of meninges, which were still connected to the vertebrae. Then, brains and spinal cords were cut into pieces and digested for 30 min at 37 °C with DNase I (Sigma-Aldrich or Roche) at a concentration of 1 mg/ml and collagenase D (Roche) at a concentration of 3.75 mg/ml. Subsequently, the tissues were mashed and passed through a 70 µm cell strainer to prepare a single-cell suspension, followed by centrifugation at 350 g for 10 min at 4 °C. The cell pellet was resuspended in 5 ml 70 % Percoll (Cytiva), which was prepared by mixing 30 ml Percoll with 18 ml Percoll Mix Solution (mixture of 90 ml 10X PBS and 264 ml H_2_O). Then, the cell suspension was slowly and carefully overlaid with 5 ml 37 % Percoll, which was prepared by diluting the 70 % Percoll solution with 1X PBS, followed by centrifugation at 650 g for 30 min at 21 °C. Then, mononuclear cells were collected from the interface between the 70 % and 37 % layer and washed twice with T cell medium to remove residual Percoll. Finally, cells were either stimulated for intracellular cytokine staining or stained directly for flowcytometric analysis.

### Production of MOG tetramer

Mouse MOG_1-125_ protein (mMOG) was produced in human embryonic kidney cells expressing the Epstein-Barr virus nuclear antigen-1 (HEK-EBNA1) according to previously published protocols (Perera et al., 2013), a cell line designed for large-scale production of recombinant proteins (Tom et al., 2008). Monomer production and tetramerization was performed as previously described (Salvador et al., 2022). Briefly, purified mMOG monomers were biotinylated using BirA ligase to enable multimerization. The biotin-binding protein streptavidin was used to form stable tetramer structures consisting of four biotinylated mMOG monomers. As the streptavidin was directly conjugated to AF 488, staining with MOG tetramer could be used to identify MOG-specific B cells by flow cytometry or confocal microscopy.

### Flow cytometry and cell sorting

For intracellular cytokine staining, cultured or isolated cells were stimulated with 50 ng/ml PMA (Sigma) and 500 ng/ml ionomycin (Sigma) in the presence of 0.7 µL/ml monensin (GolgiStop; BD Biosciences) for 4 h. For surface stainings, the cells were washed with PBS. To exclude dead cells and debris, the samples were stained with the Zombie Violet^TM^ or UV^TM^ fixable viability kit (Biolegend) diluted 1:600 in PBS for 20 min at RT. Then, cells were washed twice with FACS buffer (PBS containing 2 % FBS and 0.05 % sodium azide), and unspecific binding of antibodies to Fc receptors was blocked by incubating the cells with TruStain FcX^TM^ (Biolegend) diluted 1:50 in FACS buffer for 15 min at 4 °C. The last step was omitted when *in vitro* differentiated T cells were analyzed. Afterwards, cells were stained with fluorophore-conjugated antibodies against surface markers diluted in FACS buffer for 20-30 min at 4 °C. If biotinylated antibodies were used, the samples were washed and subsequently stained with fluorophore-conjugated streptavidin (SA) diluted in FACS buffer for 20 min at 4 °C. After two washing steps, cells were resuspended in FACS buffer for analysis. For intracellular cytokine staining, cells were fixed for 30 min at 4°C with 0.4 % paraformaldehyde (PFA, Merck KGaA) and permeabilized by washing twice with permeabilization buffer (PBS containing 2 % FBS and 0.1 % saponin (Sigma)). Then, samples were incubated with antibodies against intracellular markers diluted in permeabilization buffer for 30 minutes at 4 °C, washed twice and resuspended in FACS buffer for analysis. The following antibodies/reagents were used for flow cytometry: anti-CD19 (1D3 or 6D5), anti-CD1d (1B1), anti-CD4 (RM4-5), anti-TCR Vα3.2 (RR3-16), anti-B220 (RA3-6B2), anti-CD11b (M1/70), anti-CD21/CD35 (7E9), anti-CD11c (N418), anti-CD69 (H1.2F3), anti-CD83 (Michel-19), anti-CD40 (HM40-3), anti-IL-21R (4A9), anti-ICAM-1 (YN1/1.7.4), anti-IFNγ (XMG1.2), anti-IL-10 (JES5-16E3), anti-IL-17A (TC11-18H10.1, all Biolegend), anti-CD138 (REA104, Miltenyi), anti-GL7 (GL7), anti-IgD (11-26c.2a, all BD Pharmingen), anti-CD45 (30-F11), anti-IgM (II/41), anti-CD86 (GL1), anti-MHC-II (NIMR-4), anti-IL-17A (eBio17B7, all eBioscience), anti-TCR Vß5.1/5.2 (MR9-4, Invitrogen), PNA (Vector Laboratories). Data were acquired on a BD LSRFortessa^TM^, BD FACSverse^TM^ flow cytometer or Beckman Coulter CytoFLEX S flow cytometer and analyzed using Flowjo software (version 10.6.2; TreeStar). For sorting of B cells, surface staining of 1×10^6^ cells from spleen and cLN or 2.7-3.7×10^6^ cells from CNS samples was performed in T cell medium for 20 min at 4 °C. After two washing steps, cells were resuspended in T cell medium containing 2 mM EDTA (Sigma), filtered through a 35 µm cell strainer and kept on ice. CD45^high^ CD11b^low^ CD4^-^ CD19^+^ were sorted using a BD FACSAria™ Fusion No1.

### Immunohistochemistry (IHC) on cryosections

To increase the chances for finding eLFs in the CNS, IHC was performed using Th17-EAE mice that had a high score for several days. The mice were perfused through the left cardiac ventricle with 10 ml PBS and 10 ml 4 % PFA. Then, the brain was dissected by carefully removing the skull using forceps and scissors. After removal of the outer muscle layer, the spinal cord was dissected by snapping each vertebra open with forceps to carefully remove it from the vertebral column in one piece leaving the leptomeninges as intact as possible. Spleens and lymph nodes were also dissected as control tissue. After post-fixation in PBS containing 4 % PFA for 2 h at 4 °C, the organs were dehydrated in PBS containing 30 % sucrose for 24 h at 4 °C. Then, they were embedded in O.C.T. compound medium (Sakura or VWR Chemicals), frozen using dry ice and stored at −80 °C. Subsequently, the brain was cut into 20 µm coronal sections on a Leica CM1850 cryostat. From spinal cord, spleen and lymph nodes, 10 µm sections (longitudinal for spinal cord and spleen) were prepared and placed on slides (Superfrost Plus^TM^, Thermo Scientific) and frozen at −80 °C.

#### Giemsa staining

To screen for eLFs, every 15^th^ section of brain and spinal cord was fixed in 10 % methanol for 10 min, air-dried for 20-30 min and stained with Giemsa-stain (AppliChem), diluted 1:20 in deionized water, for 45 min. After washing with deionized water, the sections were dried overnight and mounted using Eukitt mounting medium (Sigma). Sections were screened for immune cell infiltrates by light microscopy using a Leica DM2500 microscope with DMC2900 CMOS camera.

#### Immunofluorescence staining

Following identification of eLF-containing sections via Giemsa staining, neighboring sections were stained with fluorescently labeled antibodies. After thawing, sections were fixed in cold acetone for 10 min and air-dried for 20-30 min. Then, sections were outlined with a Pap-pen and O.C.T. medium was rinsed off twice with PBS. Afterwards, sections were blocked with PBS containing 1 % bovine serum albumin (BSA) and 1:300-diluted TruStain FcX™ (Biolegend) for 1 h at RT, and incubated with primary antibodies diluted in PBS containing 1 % BSA in a chamber humidified with PBS for 1 h at RT, followed by three washing steps with PBS containing 0.1 % Tween20 for 7 min. Then, sections were stained with secondary antibodies diluted in PBS containing 1 % BSA in a dark, humidified chamber for 1 h at RT. When intracellular markers, such as activation-induced cytidine deaminase (AID), were stained, the sections were incubated with antibodies diluted in PBS containing 1 % BSA and 0.1 % saponin to permeabilize the tissue. Following three washing steps with PBS containing 0.1 % Tween20 for 5 min, sections were mounted with VECTASHIELD Antifade Mounting Medium with DAPI (Vector Laboratories) and analyzed by confocal microscopy using a Leica SP8X WLL microscope. The following antibodies/reagents were used for immunofluorescence staining: Syrian hamster anti-mouse CD3e (500A2), rat anti-mouse B220 (RA3-6B2), mouse anti-mouse/human Ki67 647 (B56, all BD Pharmingen), rabbit anti-mouse/human laminin (Abcam), rat anti-mouse/human CD11b AF 647 (M1/70), rat anti-mouse CD1d (1B1, both BioLegend), rat anti-mouse/human AID (mAID-2, eBioscience), PNA (biotinylated, Vector Laboratories), goat anti-hamster AF 568, goat anti-hamster AF 647 (both Invitrogen), goat anti-rabbit AF 568, goat anti-rabbit AF 647, goat anti-rabbit Cy3, goat anti-rat AF 633, streptavidin AF 488 (all Life technologies).

#### MOG tetramer staining

Sections were fixed in cold acetone for 5 min and O.C.T. medium was rinsed off three times with PBS for 5 min. Then, sections were outlined with a Pap-pen and blocked with PBS containing 4 % BSA, 4 % goat serum and 1:300-diluted TruStain FcX™ (Biolegend) for 1 h at RT in a humidified dark chamber. After washing three times with PBS for 5 min, sections were incubated with rat anti-mouse B220 (1:200, clone RA3-6B2, BD Pharmingen) and mMOG_tet_-SA488 (1:1000, housemade) diluted in PBS containing 4 % BSA and 1 % goat serum overnight at 4 °C. Subsequently, sections were washed three times with washing buffer (PBS containing 0.3 % Triton and 1 % goat serum) for 5 min and fixed in 4 % PFA for 30 min at RT. After three washing steps in washing buffer, sections were incubated with rabbit anti-SA488 (1:10000, Life technologies) for 3 h and goat anti-rat AF 633 (1:200, Life technologies) for 1 h diluted in PBS containing 4 % BSA and 1 % goat serum at RT in a humidified dark chamber. After three washing steps in washing buffer, sections were incubated with goat anti-rabbit Cy3 (1:1000, Life technologies) diluted in PBS containing 4 % BSA and 1 % goat serum for 3 h at RT in a humidified dark chamber, followed by three washing steps in washing buffer. Then, sections were mounted with VECTASHIELD Antifade Mounting Medium with DAPI (Vector Laboratories) and analyzed by confocal microscopy using a Leica SP8X WLL microscope.

### Enzyme-linked immunosorbent assay (ELISA)

Anti-MOG IgG1 antibodies were measured in the serum of Th17- and Th1-EAE mice by ELISA. 96 well high affinity, protein-binding plates (Nunc Maxisorp) were coated with rMOG (housemade) at a concentration of 10 µg/ml in PBS and incubated overnight at 4 °C. After washing four times with wash buffer (PBS containing 0.05 % Tween20), the plate was blocked with blocking solution (PBS containing 10 % FBS) for 1-2 h at RT, followed by four washing steps. Then, the serum samples and the antibody standard (mouse anti-MOG IgG1; clone 8.18c5; housemade) were diluted in blocking solution, added to the plate and incubated overnight at 4 °C. Following extensive washing, the plate was incubated with biotin-labeled detection antibody rat anti-mouse IgG1 (clone A85-1; BD Pharmingen) at a concentration of 1 µg/ml diluted in blocking solution for 1 h at RT. Then, the plate was washed again and incubated with avidin-horseradish peroxidase (Av-HRP; eBioscience) diluted 1:1000-1:2000 in blocking solution for 30 min at RT. After extensive washing, TMB Substrate (BioLegend) was used to induce the Av-HRP-dependent color reaction, which was stopped by addition of TMB Stop Solution (BioLegend), and the optical density (OD) was measured at 450 nm. Samples were tested in duplicates and results were reported as mean concentrations according to the standard curve.

### Single-cell RNA sequencing (scRNA-seq)

#### Library construction and sequencing

For each mouse, B cells from CNS, spleen and cLNs were pooled (90.000 B cells in total), centrifuged and, after a washing step, resuspended in PBS containing 0.04 % BSA at a concentration of 1000 cells/µl. The viability of cells was confirmed by staining with trypan blue solution. Further processing of samples was performed using the 10x Chromium Single Cell 5’ Solution (Chromium Next GEM Single Cell V(D)J v1.1 with Feature Barcoding Technology for Cell Surface Protein, 10x Genomics) according to the manufacturer’s instructions. Prior to loading on the chip, 5000 cells of each compartment were labeled with anti-mouse hashtag oligonucleotide (HTO) antibodies (TotalSeq™-C0301/C0302/C0303, Biolegend) and pooled together. Next, the B cells were processed with enzymes and gel beads, followed by partitioning in oil suspension to create gel beads in emulsion (GEMs) containing single cells. In GEMs, the cells’ RNA was reverse-transcribed into cDNA, whereby all the cDNA from the same cell was labeled with a shared barcode. scRNA-seq, scBCR-seq and cell hashing libraries were multiplexed using individual Chromium i7 Sample Indices. Quality controls were performed with Qubit dsDNA HS kit (Life Technologies) to quantify the libraries, and Bioanalyzer 2100 (Agilent Technologies) to confirm fragment length. Then, the resulting cDNA libraries were pooled and send for sequencing. scRNA-seq and scBCR-seq libraries were sequenced on NovaSeq S4 Flowcells using 150-bp paired-end reads and 8LJbp for the i7 index, aiming for 50,000 reads per cell for gene expression and 5,000 reads per cell for BCR enrichment. Cell hashing libraries were sequenced on NovaSeq S1 Flowcells using 50-bp paired-end reads and 8LJbp for the i7 index, aiming for 10,000 reads per cell.

#### Data processing

To demultiplex samples, align the reads to the reference mouse mm10 genome, and process the raw data, we used the Cell Ranger Software (10X Genomics, v. 3.1.0). Following this, the unique molecule identifiers (UMIs) were summarized and the single-cells were filtered by the number of UMI counts identified. Then, we processed the gene matrices with R (v. 4.0.5) and the R package Seurat (v. 4.0.6 and 4.9.9) (Butler et al., 2018; Stuart et al., 2019). Cell hashing raw counts were normalized using centered log-ratio (CLR) transformation by comparing the counts of single cells to the geometric mean of the HTOs signal with subsequent logarithmic transformation. Cells with missing HTO signals or containing two different HTO-identifiers were removed from the further analysis. Further, the scRNA-seq count matrices were log-normalized and scaled by passing through only the cells containing more than 200 and less than 5000 genes with mitochondrial gene fractions below 5 %. Subsequently, the highly variable genes were identified with the vst method, and cells from each compartment were integrated using the canonical correlation analysis (CCA) method. The aligned BCR sequencing information was added into the metadata file of the Seurat object with subsequent removal of cells with missing BCR sequences. The cells were considered as expanded if at least 2 cells belonged to the same B cell clonotype. In the next step, 18413 cells were recovered and the corresponding PCA was computed using the previously identified highly variable genes. Based on the first 15 PCA dimensions, we constructed the Shared Nearest Neighbor (SNN) graph and clustered the cells by using the FindNeighbours and FindClusters functions of the Seurat R package. The following dimensional reduction was performed with the help of the Uniform Manifold Approximation and Projection (UMAP) method. One cluster, cluster 11, with markedly lower quality compared to the other clusters based on nFeature and nCount, was removed from further analysis. In addition, cluster 10 was identified as expressing macrophage/microglia markers and was therefore also excluded (based on AddModuleScore from Immgen Monocytes MHCIIhi blood vs Fo B cells spleen and Microglia CNS vs FoB form spleen).

For identification of differentially upregulated genes per cluster, compartment, or expansion condition we used the FindMarkers function based on the log-transformed count matrices of each single cell by applying the Wilcoxon test and subsequent p-value adjustment with the Bonferroni correction, with logfc.threshold = 0.

#### Pathway network analysis

For each cluster, GSEA (Mootha et al., 2003; Subramanian et al., 2005) PreRanked files were generated with the gene and avg_log2FC columns. GSEA v4.2.3 was run with the mouse GO BP database m5.go.bp.v2023.1.Mm.symbols (https://data.broadinstitute.org/gsea-msigdb/msigdb/release/2023.1.Mm/), without collapsing gene symbols, scoring_scheme classic, set_max 500, set_min 2, and otherwise default parameters. The results were exported to Cytoscape v3.9.1 (Shannon et al., 2003), where only the nodes with NES > 0, fdr_qvalue < 0.1 and fwer_qvalue < 0.5 were kept. AutoAnnotate from the EnrichmentMap app collection (Merico et al., 2010) was used to cluster and label the pathway networks, with Cluster Algorithm = MCL cluster, Label Column = Node Names, Max Words Per Label = 4, Minimum Word Occurrence = 1, Adjacent Word Bonus = 8, setting numbers and “GOBP” as extra Excluded Words.

#### B cell functional signatures

Functional signatures expressed by B cells were calculated with the AddModuleScore function from Seurat from manually curated gene lists based on ImmGen (https://www.immgen.org/), GO terms (https://www.informatics.jax.org/vocab/gene_ontology) and literature search.

<u>APC/synapse signature:</u> this signature was assembled by combining the GO gene set “lymphocyte co-stimulation” (GO:0031294, without T cell specific genes) and the GO gene set “immunological synapse” (GO:0001772, without T cell specific genes), and then adding the core APC signature described by (Jordão et al., 2019), the list of molecules described to be present in the immunological synapse formed by primary murine B cells by Cunha et al. (https://www.biorxiv.org/content/10.1101/2023.02.23.529674v1.full.pdf), and manually adding Icosl, Tnfsf9, Timd4, Pvr, and Il21r.

<u>B1a signature:</u> this gene set is derived from Immgen by population comparison using the ULI RNAsequencing datasets. B1a cells from spleen and B1a cells from the peritoneal cavity were compared against follicular B cells from spleen, and Nfatc1 and Mzb1 were added manually.

<u>MZ signature:</u> this gene set is derived from Immgen by population comparison using the ULI RNAsequencing datasets. Here, marginal zone B cells from spleen were compared to follicular B cells from spleen and Nfatc1 and Mzb1 were added manually.

<u>LZ-GC and DZ-GC signature:</u> these signatures were described by (Victora *et al*., 2010).

<u>mTOR signaling:</u> Hallmark_mtorc1_signaling (mouse) gene set from the MSigDB database.

<u>Myc signaling:</u> Hallmark_myc_targets_v1 (mouse) gene set from the MSigDB database.

<u>OXPHOS:</u> Hallmark_oxidative phosphorylation (mouse) gene set from the MSigDB database.

<u>PB signature:</u> this gene set is derived from Immgen by population comparison using the ULI RNAsequencing datasets. Here, plasmablasts and plasma cells from spleen and plasma cells from bone marrow were compared against follicular B cells from spleen.

<u>IFNα response:</u> Hallmark_interferon_alpha_response (mouse) gene set from the MSigDB database.

<u>IFNγ response:</u> Hallmark_interferon_gamma_response (mouse) gene set from the MSigDB database.

<u>TNFα signaling:</u> Hallmark_TNFa_signaling_via_nfkb (mouse) gene set from the MSigDB database.

<u>IL-21 response:</u> this signature is described on Immgen and includes all genes upregulated in follicular B cells at least 2-fold by IL-21 relative to PBS treatment.

<u>B cell activation:</u> this signature represents the GO gene set “B cell activation involved in immune response” (GO:0002312).

### Data and code availability

All sequencing data (fastq files, processed data files and Seurat Object) from this study is available upon request. Custom code used for analysis is available at Zenodo under the same doi as the published manuscript.

### Intravital two-photon microscopy

#### Animal preparation

Th17 recipients, which had been on a high score for several days to allow for continued immune cell infiltration and maturation of eLFs, were selected for live imaging analysis. Prior to surgery, the mice were anesthetized with MMF anesthesia consisting of midazolam (5 mg/kg bodyweight (bw)), medetomidine (500 µg/kg bw) and fentanyl (50 µg/kg bw). Subsequently, they were tracheotomized and ventilated with 1.5-2 % isoflurane, which was supplied throughout the surgery and the entire imaging session. Thereby, the mice were placed on a custom-made microscope stage, while their body temperature was stabilized at 37.5 °C using a heated pad. During imaging, breathing parameters, including concentrations of inspiratory and expiratory gases as well as ventilation pressure, and electrocardiograms were constantly monitored. The experimental procedure of spinal cord imaging has been described previously (Bartholomäus et al., 2009; Mues et al., 2013)). In brief, a small imaging window at level Th12/L1 of the spinal cord was prepared. First, a midline incision of about 2 cm was placed to open the skin and, subsequently, to detach the paravertebral musculature. To reduce artefacts caused by breathing and thereby ensure stable imaging conditions, a custom-made stage was used in order to fix three spines. Then, a laminectomy on the central vertebra was performed using a dental drill (Foredom). Finally, low-melting agarose was used to build a ring-shaped surrounding, which was filled with PBS to keep the tissue of the opened spine hydrated and to allow using the water immersion objective.

#### Image acquisition

Time-lapse two-photon laser-scanning microscopy was performed using a SP8 multi-photon microscope (Leica). The SP8 system was equipped with a pulsed InSight DS+ laser (Spectra Physics). The emission wavelength was tuned to 841 nm and the fluorescent signal was collected using a water immersion objective (25x, NA 1, Leica) and detected with external, non-descanned hybrid photo detectors (HyDs) featuring 483/32 nm, 535/30 nm, 585/40 nm and 650/50 nm bandpass filters. Images were recorded using a 1-2x zoom, whereby areas of 221x221 µm, 295x295 µm or 443x443 µm were scanned, and 49-115 µm z-stacks with 2.5-4.3 µm step size were acquired with an acquisition rate of 21-22 s per z-stack.

#### Image processing and analysis

Time-lapse images were acquired using the LAS X software (Leica), followed by processing and analysis using ImageJ (NIH). Before the stacks were z-projected with maximum intensity to obtain two-dimensional movies, a Gaussian blur filter was applied. For Twitch-2B-expressing cells, the cpVenus^CD^ Fluorescence resonance energy transfer (FRET) channel was divided by the mCerulean3 (CFP) channel and, by changing to a fire lookup table, ratiometric pseudocolor images were generated. In each time frame of the two-dimensional maximum intensity projection, the cell shapes were outlined manually to create a region of interest (ROI) for analyzing the calcium ratios. Since the bleed-through of CFP into the YFP channel was determined to account for 44 %, the FRET signal was corrected as follows: YFP= FRET – 0.44 x CFP. In order to calculate the calcium ratio in every time frame, the average signal intensities of all pixels in each ROI were used. Cell coordinates over time were used to calculate motility parameters by using ImageJ. T:B cell contact analysis was performed by monitoring the movement of each Twitch-2B^+^ T cell and its interaction with tdTomato^+^ B cells in the 3D follicle structure using the LAS X software.

### Statistical analysis

Statistical analysis was performed using GraphPad Prism software (version 7). The statistical test chosen is described in the figure captions. For comparison of two groups, statistical differences were calculated using the unpaired Student’s *t*-test for normally distributed data, including Welch’s correction in case of unequal variances, or the Mann-Whitney U test if Gaussian distribution could not be determined. For comparison of three groups, the one-way ANOVA with Tukey’s test for multiple comparisons was applied.

## Acknowledgements

We thank the Core Facility Flow Cytometry and the Core facility Bioimaging at the Biomedical Center, Ludwig-Maximilians University Munich, for providing equipment, services and expertise. We are grateful to the staff from CCGA Kiel for their assistance with sequencing and data transfer. We thank Isabel Gök and Klaus Hämäläinen for technical assistance. The study was supported by the German Research Foundation (DFG) through the Emmy Noether programme PE-2681/1–1 (to A.P.), through SFB TRR 128 (project B08 to A.P.; project B10 to N.K. and M.K.), through CRC TRR 152 (ID 239283807; project P27; to A.P. and M.K.), through the Heisenberg programme (KA2951/3-1 and KA2951/3-2 (264119140) to N.K. and through research grant KA2951/2-2 (246754395) to N.K.

