## Supplementary material for "T:B cell cooperation in ectopic lymphoid follicles propagates CNS autoimmunity": Supplemetary Figure legends

**Supplementary Material to the Manuscript “T:B cell communication in ectopic lymphoid follicles in CNS autoimmunity”**

Figure S1. Th1 and Th17 adoptive transfer EAE.

Naïve MOG-specific T cells isolated from 2D2 mice are differentiated with polarizing cytokines into Th1 or Th17 cells in the presence of anti-CD3 and APCs. After 5-8 days, the cells are restimulated with anti-CD3 and anti-CD28 for 2 days before they are transferred i.v. or i.p. into recipient animals. Clinical disease develops after 8-15 days post transfer. (**A**) After 4 days under Th1 or Th17 polarizing conditions, cultures were analyzed for IL-17A and IFNγ expression by intracellular cytokine staining. Representative flow cytometry plots are shown. (**B**) EAE incidence (upper panel) and mean EAE scores (lower panel) of Th1 and Th17 recipients that developed clinical disease. Mean ± SEM. Clinical data of one experiment (n_Th1_ = 4 mice; n_Th17_ = 8 mice) is shown. Similar results were obtained in four (Th1-EAE) or 14 (Th17-EAE) independent experiments. (**C**, **D**) At the peak of disease, cells were isolated from the CNS of (**C**) Th1 and (**D**) Th17-EAE mice and analyzed by flow cytometry. Transferred T cells were identified by their expression of the transgenic MOG-specific Vα3.2 TCR. Transferred and endogenous (Vα3.2^-^) T cells were analyzed for expression of IL-17A and IFNγ by intracellular cytokine staining. Flow cytometry plots are representative of (**C**) three independent experiments with two to three mice or (**D**) seven independent experiments with two to five mice.

**Figure S2. Large eLFs form in the spinal cord and brain of Th17-EAE mice.** At the peak of disease, (**A**, **B**, **D**) longitudinal spinal cord and (**C**) coronal brain cryosections of Th17-EAE mice were prepared. (**A**) Every 15^th^ section was stained with Giemsa stain to identify regions with lymphocytic infiltrates. (**B**, **C**) Cryosections were stained for T cells (CD3, red), B cells (B220, green) and laminin (blue). (**C**) Subarachnoid space at the base of the brain. (**D**) Cryosections were stained for T cells (CD3, red), B cells (B220, green) and macrophages (CD11b, cyan) Scale bar: (A, D) 100 µm, (B, C) 200 µm.

**Figure S3. T and B cells express germinal center markers in the CNS of Th17-EAE mice.** CNS-infiltrating cells isolated at the peak of disease were investigated by flow cytometry. B cells were analyzed for expression of (**A**) PNA, (**B**) IgM and IgD (pre-gated on CD19^+^ cells) and (**C**) the plasma cell marker CD138 (pre-gated on CD45^high^ CD11b^low^ cells). (**D**) Transferred (pre-gated on Vα3.2^+^ CD4^+^) and endogenous (pre-gated on Vα3.2^-^ CD4^+^) T cells were analyzed for expression of the germinal center marker GL7. Flow cytometry plots are representative of (**A**) 7 mice from two independent experiments, (**B**) 17 mice from four independent experiments, (**C**) 9 mice from two independent experiments and (**D**) 17 mice from five independent experiments.

**Figure S4. Comparative gene expression analysis of all clusters.** At the peak of disease, B cells were sorted from indicated organs of Th17 recipients (n = 3 mice) based on their expression of CD19, and their gene expression was characterized by single-cell RNA sequencing. (**A**) Heatmap of gene expression levels of cluster 0-9. (**B**) Proportion of cells belonging to cluster 0-9 in (upper panel) CNS, (middle panel) spleen and (lower panel) cLN. (**C**) Mean module score of gene expression associated with (left panel) *Mtor* signaling, (middle panel) *Myc* signaling and (right panel) oxidative phosphorylation *OXPHOS* in (colored) the CNS-enriched clusters 6-9 and (grey) the remaining clusters.

**Figure S5. Properties of CNS-specific clusters.** (**A**-**C**) At the peak of disease, B cells were sorted from indicated organs of Th17 recipients (n = 3 mice) based on their expression of CD19, and their gene expression was characterized by single-cell RNA sequencing. Pathway network analysis of upregulated pathways in (**A**) cluster 6, (**B**) cluster 7 and (**C**) cluster 9. (**D**, **E**) Cells were isolated from the CNS of Th17 recipients and analyzed for expression of CD21 and CD1d by flow cytometry. A pre-gate on CD19^+^ IgM^high^ IgD^low^ B cells was applied. (**D**) Representative plot and (**E**) quantification of the frequency of CD21^+^ CD1d^+^ cells of 8 mice from two independent experiments. (**F**) Spinal cord cryosections of Th17 recipients were stained for T cells (CD3, blue), B cells (B220, red) and CD1d (green). Scale bar: 50 µm. (**G**-**J**) At the peak of disease, B cells were sorted from CNS, spleen and cLN of Th17 recipients (n = 3 mice) based on their expression of CD19, and characterized by BCR repertoire analysis. (**G**) Feature plot depicting distribution of nonexpanded (grey) and expanded (colored) B cells among B cell clusters (all organs). The color code indicates the number of cells per clone. (**H**) Zoom of cluster 8 from (**G**). (**I**) Pathway network analysis of upregulated pathways in cluster 8. (**J**) Violin plots showing expression level of *Fam46c*, *Emb* and *Gpx4* in nonexpanded and expanded cells (with expansion factors ≥3) in cluster 8.

Figure S6. Transferred T cells recruit endogenous MOG-specific B cells. (A) MOG-specific IgG1 antibodies were measured via ELISA in the serum of Th17 and Th1 recipients at different time points (days) after adoptive transfer. Dots represent individual mice. Mean ± SEM, Mann-Whitney U test (Th17 pre-onset vs. peak of disease), Welch`s t-test (Th1 early disease vs. peak of disease). *p < 0.05; **p < 0.01. (B) As positive control, lymph node cryosections from an IgH^MOG^ mouse were stained for B cells (B220, green) and MOG tetramer (red). Scale bar: 100 µm. (C) CNS cryosections from Th17-EAE mice at the peak of disease were stained for B cells (B220, green) and MOG tetramer (red). (Left panel) While most eLFs contain only very few MOG tetramer^+^ B cells (arrow), (right panel) some eLFs show massive accumulation of MOG tetramer^+^ B cells. Box in left lower corner shows zoom of MOG tetramer staining. Scale bars: 100 µm.

**Figure S7. CNS-infiltrating B cells show elevated expression of genes associated with B cell activation, co-stimulation and antigen presentation/immune synapse.** (**A**) Violin plots showing expression levels of genes induced in response to IFNα. Two-sided *t*-test with FDR adjusted p-values. *p < 0.05; **p < 0.01; ***p < 0.001, ****p < 0.0001. Significantly downregulated gene sets are marked in grey. (**B, C**) At the peak of disease, cells were isolated from CNS or spleen and B cells were analyzed for expression of (**B**) CD69, CD83, CD86 and (**C**) MHC-II, CD40, ICAM-1, IL-21R (pre-gated on CD19^+^ cells). Flow cytometry plots are representative of (**B**) (CD69) 5 mice or (CD83 and CD86) 6 mice and (**C**) 6 mice.

Figure S8. Studying interactions of Th17 cells expressing the calcium indicator Twitch-2B and B cells expressing tdTomato in meningeal eLFs by intravital microscopy. (A) During *in vitro* differentiation, T cells were either transduced after two days in Th17 conditions (primary phase) or one day after restimulation (secondary phase) and tested for transduction efficiency after two days. Flow cytometry plots are representative of four independent experiments. (B) The frequency of Twitch-2B^+^ T cells before adoptive transfer (secondary phase) and in the CNS of Th17-EAE mice was compared. For the CNS sample, a pre-gate on transferred (Vα3.2^+^) T cells was applied. Flow cytometry plots are representative of nine mice from four independent experiments. (C) Twitch-2B^-^ and Twitch-2B^+^ T cells from the CNS of Th17-EAE were analyzed for expression of IL-17A and IFNγ by intracellular cytokine staining. A pre-gate on transferred (Vα3.2^+^) T cells was applied. Flow cytometry plots are representative of four mice from two independent experiments. (D) CNS CD19^+^ B cells from Mb1.Cre x ROSA26 tdTomato Th17-EAE mice were analyzed for tdTomato expression by flow cytometry (left) and confocal microscopy of a CNS cryosection stained for T cells (CD3, blue) and B cells (B220, green) (right). The shown flow cytometry plot is representative of six mice from three independent experiments. Scale bar: 50 µm. (E) Duration of calcium signaling (min) of 2D2 and OT-II T cells. The dotted line at 2 min indicates the cut-off between short-term and long-term calcium signaling. Mann-Whitney U test. **p < 0.01. (E, F) Representative tracks of (F) 2D2 T cells (n = 104 cells from three independent experiments) and (G) OT-II T cells (n = 107 cells from four independent experiments) showing calcium indicator ratio (YFP/CFP, red) and velocity (black) in each time frame. The red dashed line indicates the ratio threshold (0.534). Grey shading highlights time frames with B cell contact.

**Table S1. Top 50 upregulated genes per cluster.**

Movie 1. Visualization of calcium signaling in T cells in meningeal eLFs. Intravital microscopy of Twitch-2B^+^ T cells and tdTomato^+^ B cells in spinal cord eLFs of Th17-EAE mice. (Left) Fluorescence overlay of Twitch-2B^+^ T cells (green) and tdTomato^+^ B cells (red) and (right) pseudocolor calcium ratio image showing calcium levels of T cells.
