## Supplementary figures and images for "T:B cell cooperation in ectopic lymphoid follicles propagates CNS autoimmunity"

### Supplementary Figure 1

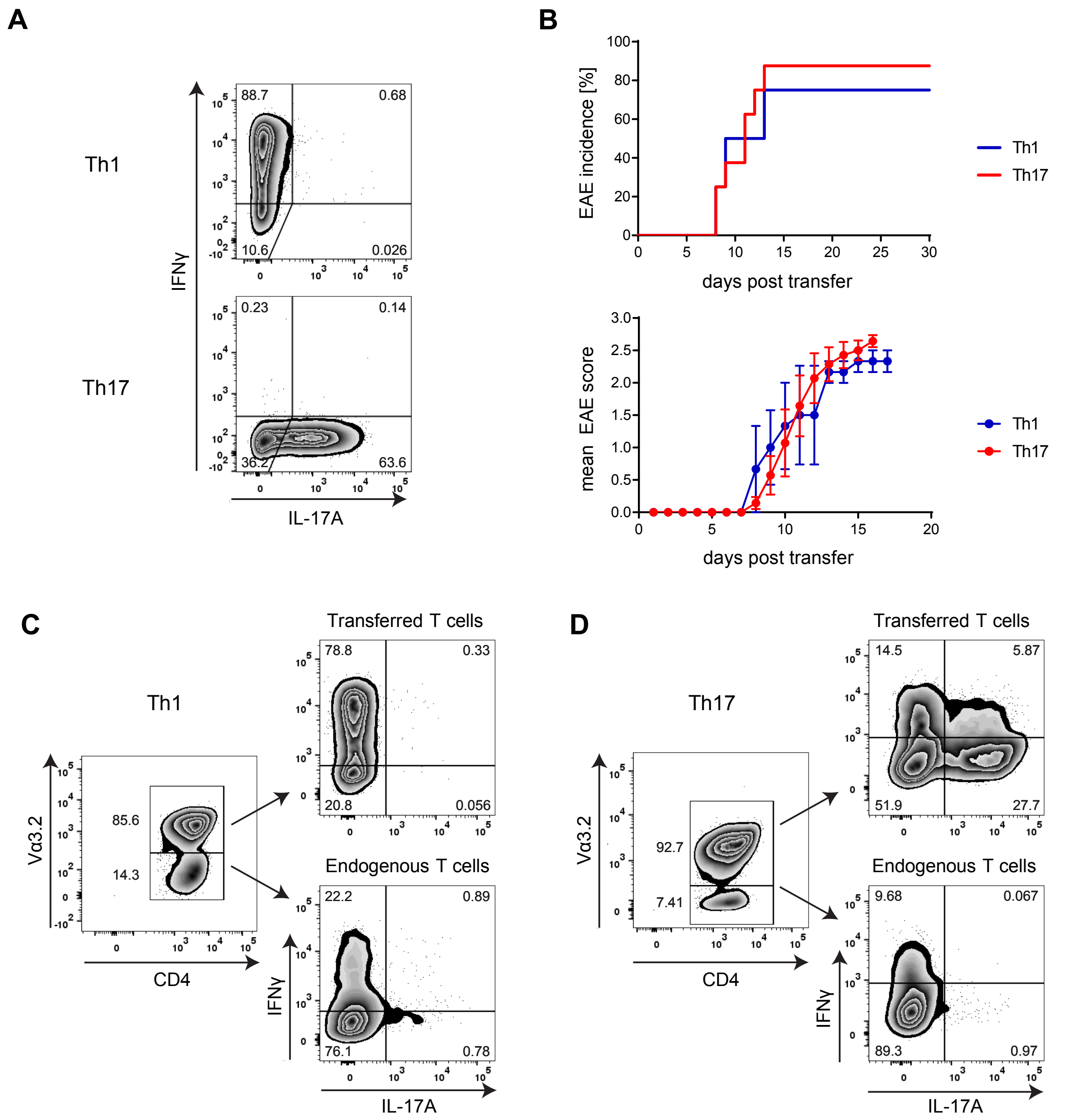

### Supplementary Figure 2

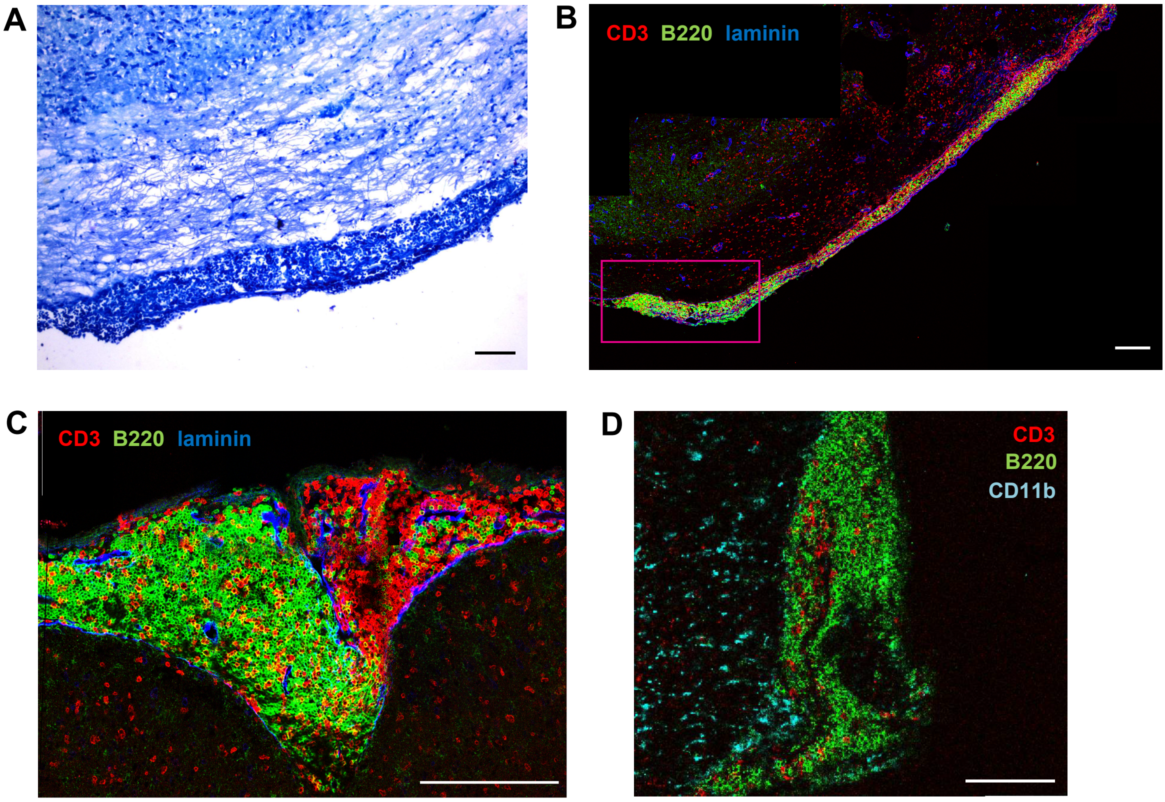


Figure S2

### Supplementary Figure 3

CNS B cells

CNS T cells

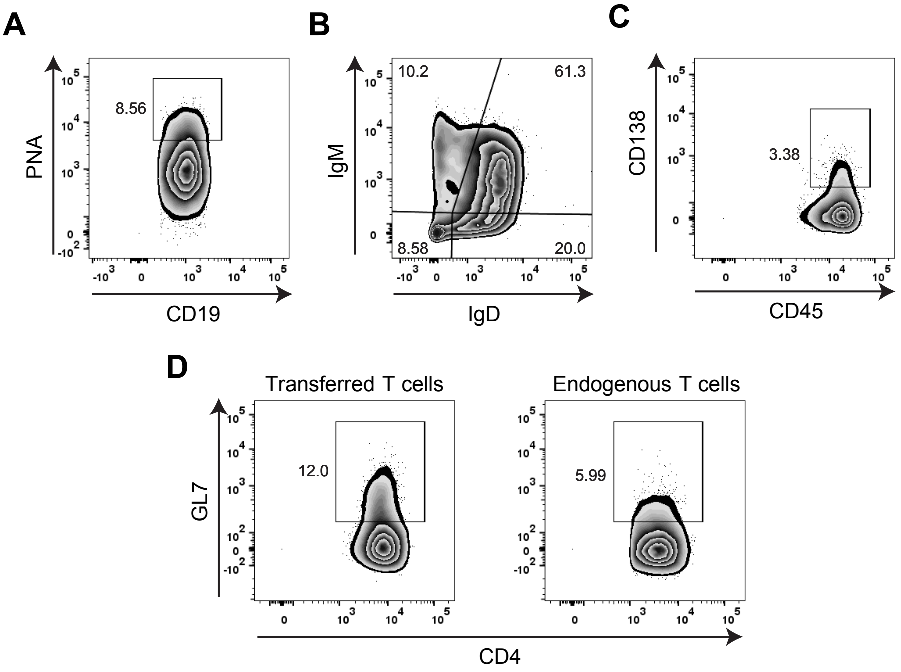


Figure S3

### Supplementary Figure 4

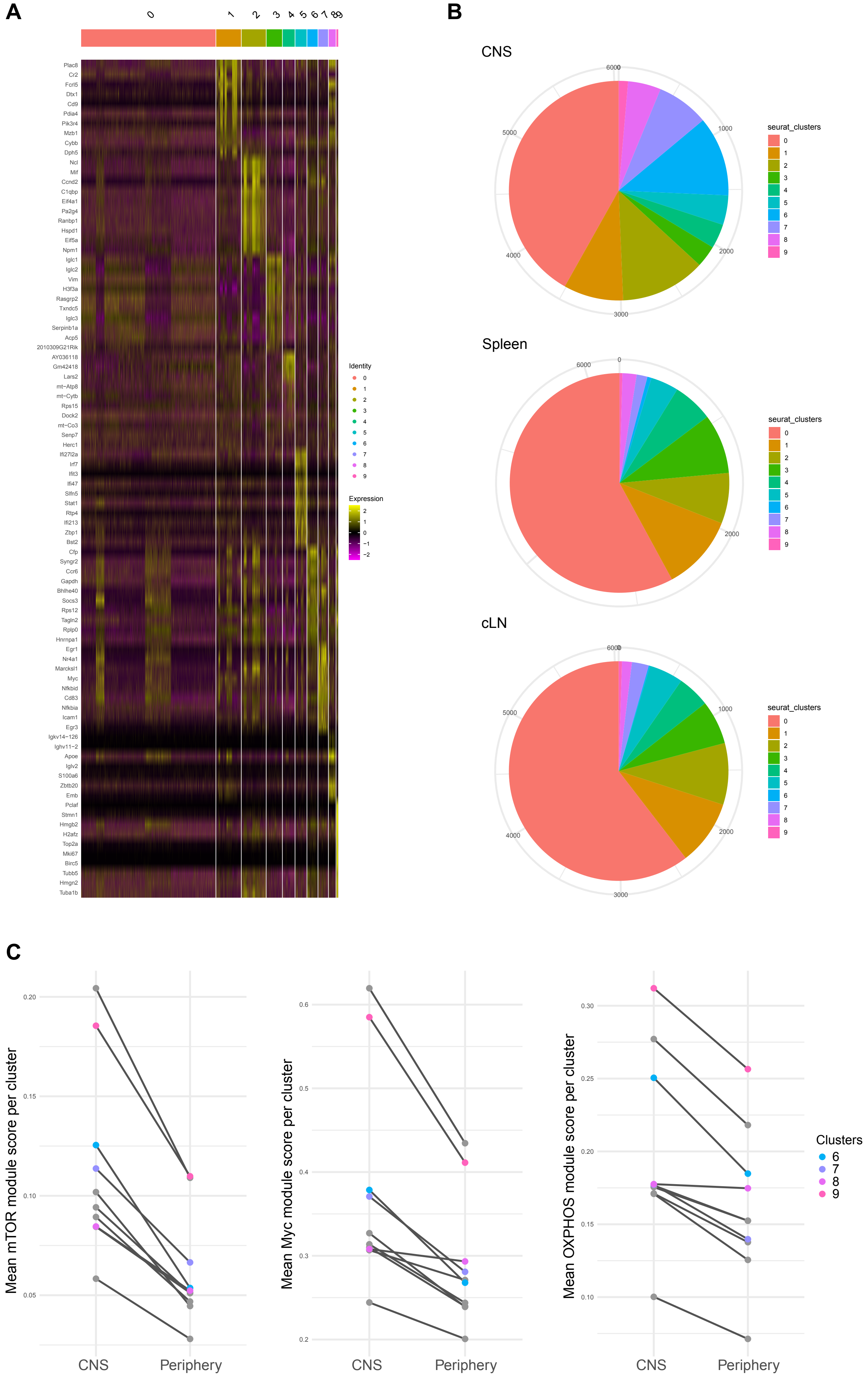

### Supplementary Figure 5

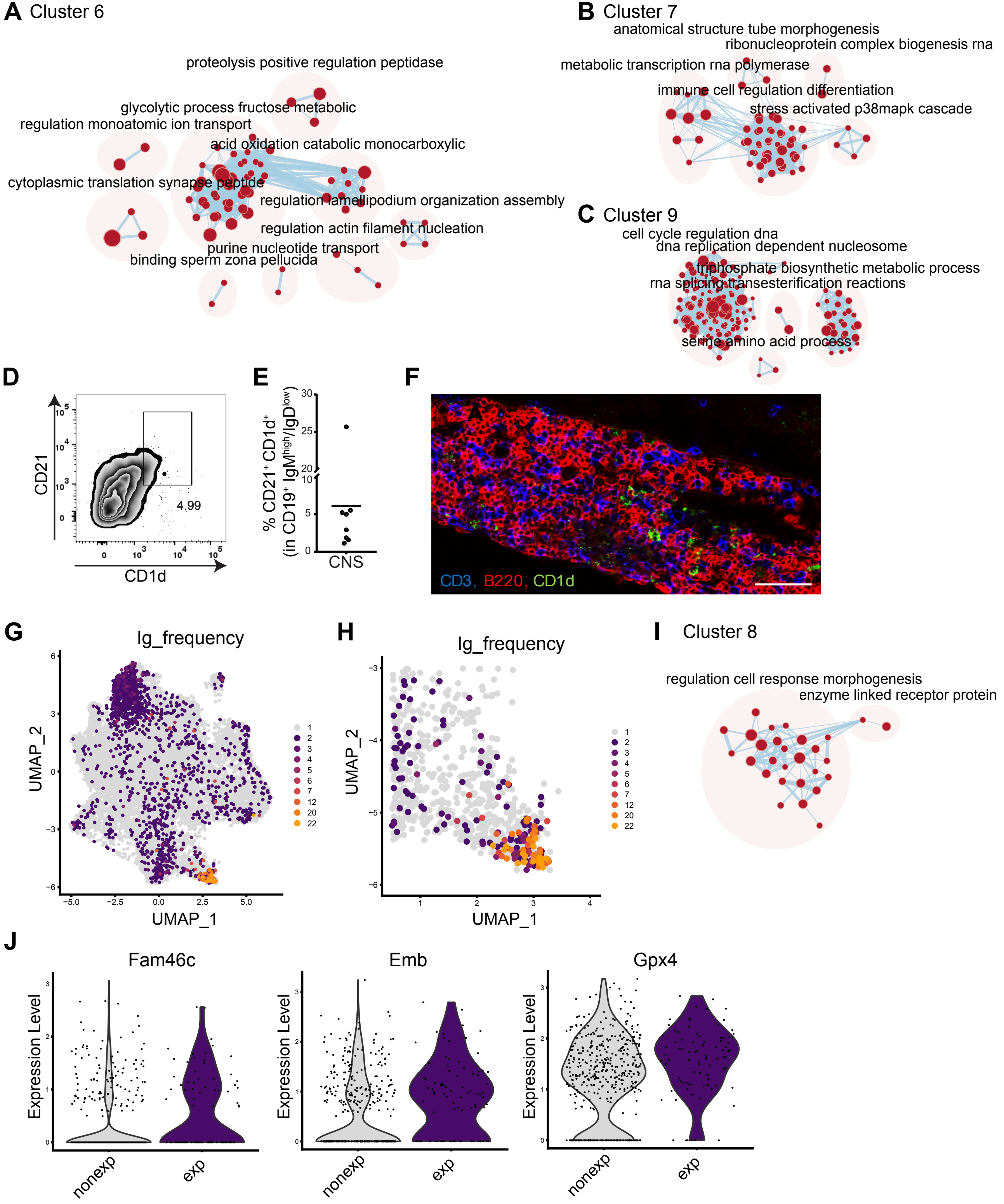

### Supplementary Figure 6

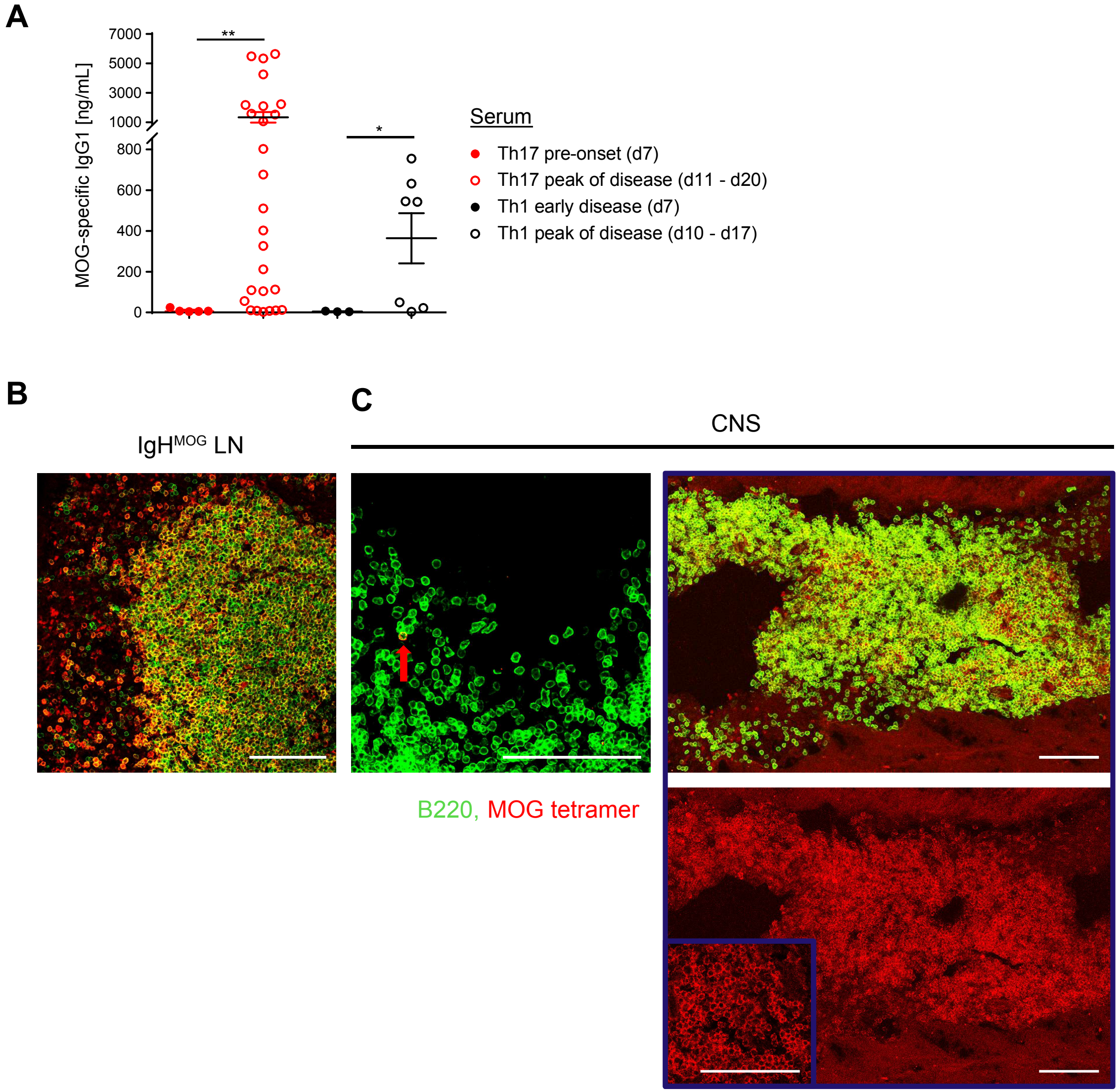

### Supplementary Figure 7

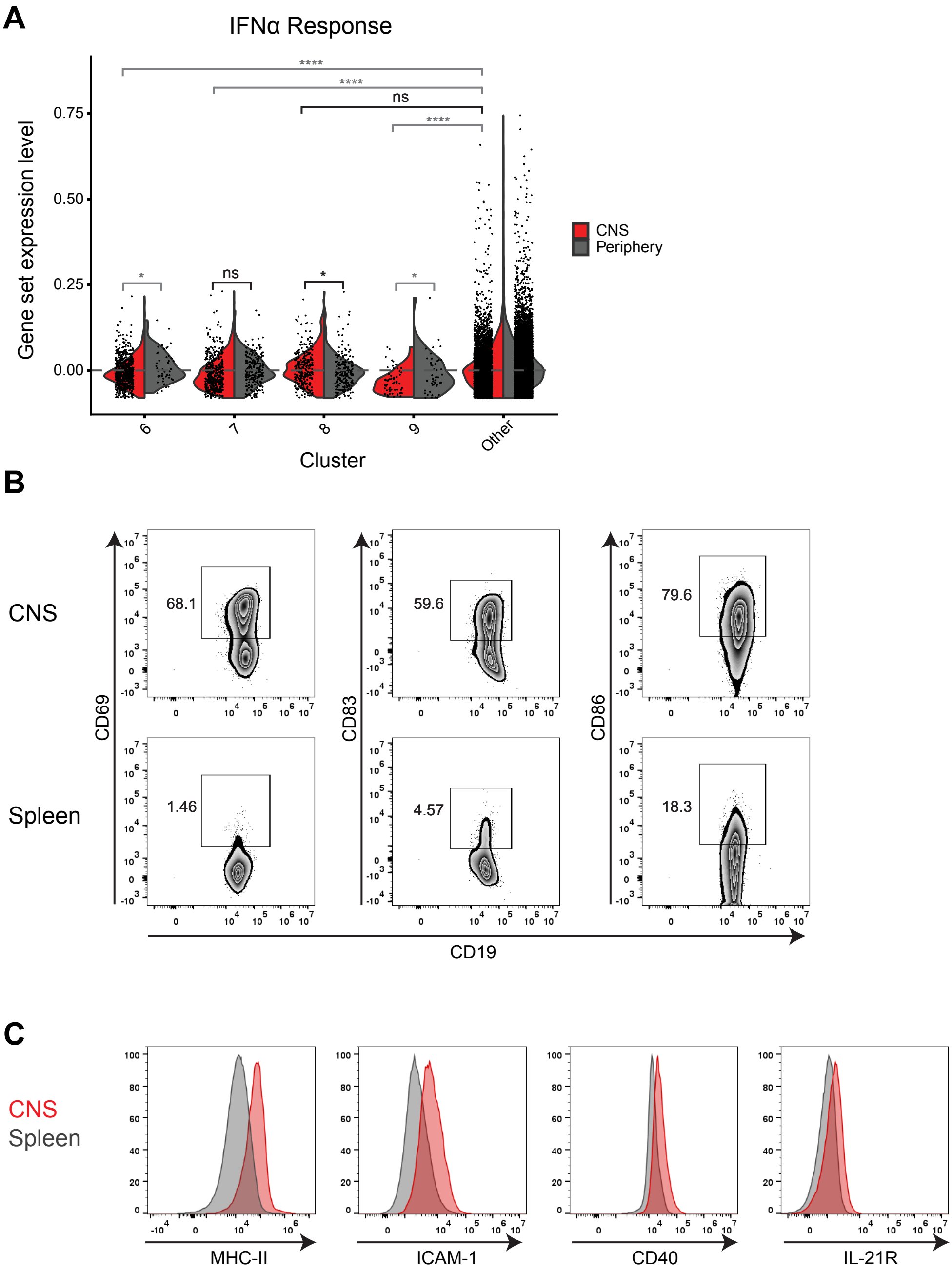

### Supplementary Figure 8

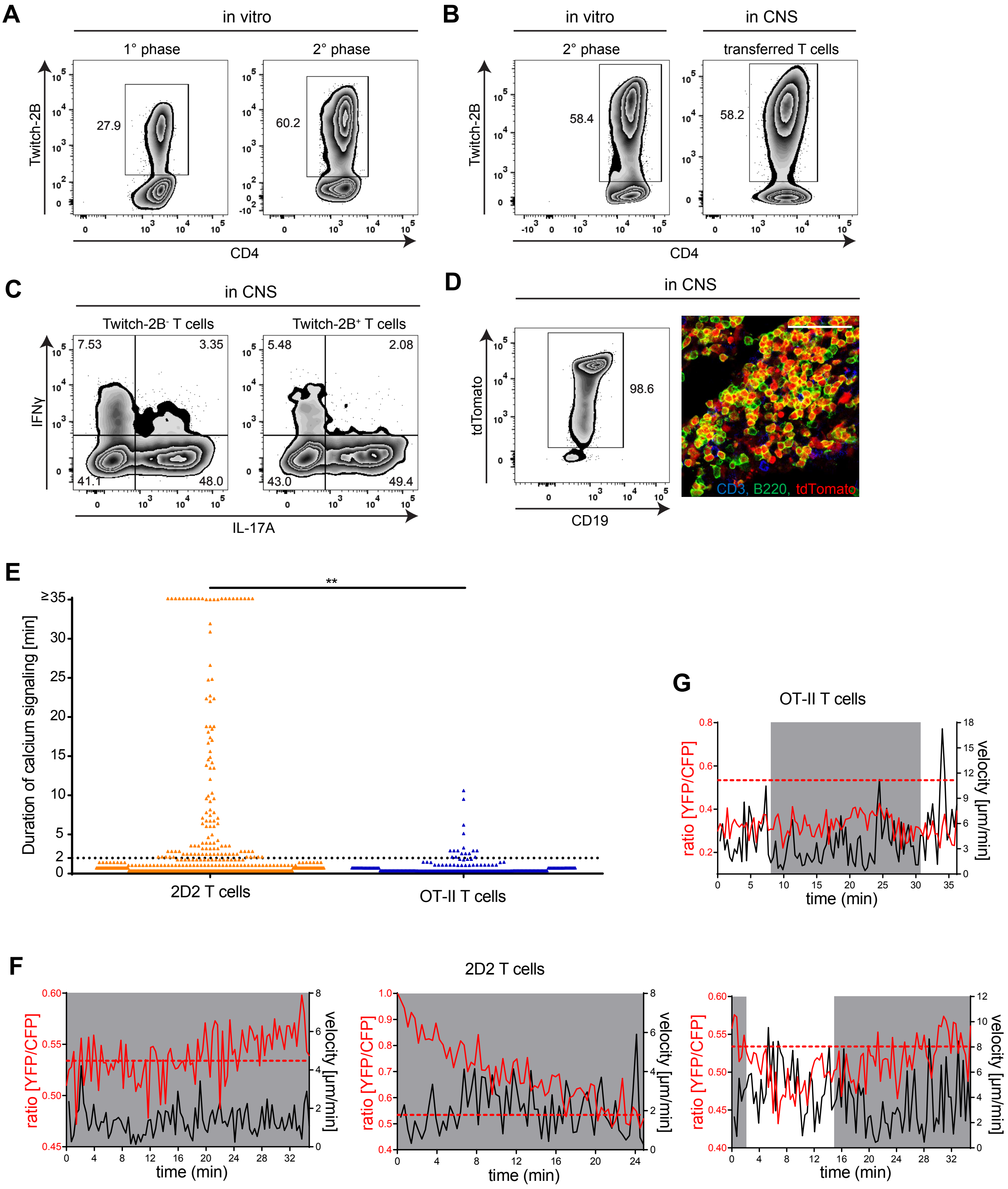
